# Fine-tuning cell-mimicking polyacrylamide microgels: Sensitivity to microscale reaction conditions in droplet microfluidics

**DOI:** 10.1101/2025.07.24.666603

**Authors:** Ruchi Goswami, Kyoohyun Kim, Aldo R. Boccaccini, Jochen Guck, Salvatore Girardo

**Affiliations:** Max Planck Institute for the Science of Light, Erlangen, Germany; Max-Planck-Zentrum für Physik und Medizin, Erlangen, Germany; Institute of Biomaterials, University of Erlangen-Nürnberg, Erlangen, Germany; Department of Physics, Friedrich-Alexander-Universität Erlangen-Nürnberg, Erlangen, Germany

**Keywords:** Polyacrylamide, free-radical polymerization, microgels, droplet microfluidics, mechanobiology, biomaterials

## Abstract

The ability to shape polyacrylamide (PAAm) hydrogels using droplet microfluidics enables the production of microgel particles that mimic the cellular physical properties, opening new avenues in mechanobiology. Precisely controlling microgel size and elasticity is crucial yet complex, as various factors influence polymerization and the resulting network structure. While it is well established that chemical reactions in microdroplets are typically faster and more homogeneous than in bulk systems, an often-overlooked aspect is the increased sensitivity of these microreactors: minor variations in chemical or physical parameters can lead to significant changes in the resulting microgel properties. Our study identifies flow conditions as a key factor influencing both microgel elasticity and size, by modulating interfacial transport during gelation. Using a flow-focusing microfluidic chip, we generated pre-gel droplets with identical composition dispersed in an oil phase, systematically varying the PAAm-to-oil flow rate ratio while keeping the total flow rate constant. This approach yields droplets with minimal diameter variation (<1 µm), yet produces beads with distinct Young’s moduli, despite identical total monomer concentrations. Further analysis revealed that the catalyst transport across the oil–water interface significantly affects the polymerization efficiency and polymer network. Our findings highlight that despite the advantages of droplet-based polymerization, achieving reproducible microgel properties requires careful control of flow parameters. This underscores the importance of precise microfluidic control in advancing PAAm microgel applications in biophysics.

## Introduction

Hydrogels are three-dimensional (3D) networks of water-insoluble yet hydrophilic polymer chains [1]. The precise tuning of their physical properties has revolutionized the field of biomaterials, allowing researchers to develop materials that closely mimic natural tissues. This advancement has paved the way for groundbreaking studies in cell-material interactions [2,3], controlled drug delivery [4] and tissue engineering [5], enabling scientists to explore fundamental questions that were previously out of reach. Polyacrylamide (PAAm) hydrogels, in particular, have become popular as model soft material in mechanobiology, especially as 2D substrates for studying cell-substrate interactions in vitro [6,7]. These hydrogels, typically formed through the radical polymerization of monomers and crosslinkers, offer tunable elasticity through modulation of the pre-gel composition [8]. Additionally, their exceptional mechanical compliance, biocompatibility and moldability into custom shapes allow the replication of physiological microenvironments, enabling the exploration of fundamental biological processes and advancement of biomaterial innovation [7]. For instance, in their pioneering study, Engler et al. showed that mesenchymal stem cell spreading and differentiation are governed by the substrate stiffness: soft substrate (∼0.1–1 kPa) induced neurogenesis while the stiff substrates (>30 kPa) led to osteogenesis [3]. Additionally, substrate stiffness and topographical cues have been shown to regulate cell migration, alignment, focal adhesion dynamics and tissue morphogenesis [9–12].

While these studies on 2D substrates have provided valuable insights, the recent development and application of PAAm microscale particles, commonly known as microgels or microgel beads [13–15], have expanded the mechanobiology toolkit by offering unique three-dimensional, deformable models. These microgels have been used as mechanical calibration standards for advanced mechanical phenotyping techniques [16–19]. As mechanosensors, they have enabled measurement of cell-scale stress in vivo [20,21] and the measurement of axial and traction forces in vitro [22–24]. When used as simplified cell models into zebrafish vasculature, they have provided insights into understanding the influence of mechanics of circulating tumor cells on their circulation and extravasation during metastasis [25,26]. Additionally, when densely packed into 3D scaffolds to form granular hydrogels, they have offered a unique ability to decouple gel porosity from elasticity, creating dynamic environments for studying cell proliferation and migration [27,28] and offering a possibility for 3D bioprinting of complex structures [29]. These applications highlight their value in probing complex cellular behaviors in physiologically relevant settings.

To fully exploit the potential of PAAm microgels and to improve their reliability in biophysical studies, precise control over microgel size and elasticity is essential. For instance, their reliable use as simplified cell models and mechanosensors depends on high monodispersity in size and tightly controlled and well-characterized mechanical properties, as the accuracy of the forces derived from analytical models is highly sensitive to bead size and elasticity [20,25,26,30]. Such control relies on the combination of microfabrication methods [13,23,31–33] with PAAm polymer chemistry, which is extensively developed and studied in bulk gels [8,34], to simultaneously control the shape and mechanical properties of the resulting microgels. The method relies on the formation of spherical microreactors that serve as templates for microgel polymerization. Among the various microfabrication techniques, droplet microfluidics stands out for its ability to generate uniform, monodisperse pre-gel droplets dispersed in a continuous oil phase with high throughput and precise control over their size and composition. The pre-gel droplet composition, in turn, dictates the final physical properties of the resulting microgels. Droplet microfluidics also offers distinct physicochemical advantages: small fluid volumes yield high surface-to-volume ratios and heat transfer [35], while laminar flows at low Reynolds numbers ensure predictable mixing occurring primarily via diffusion rather than turbulence [36]. These features support homogeneous reactions, reduce concentration gradients and enhance reproducibility [37,38]. As a result, polymerization proceeds more rapidly and uniformly than in bulk gel, making droplet microfluidics an ideal platform for producing microgels tailored for biophysical applications.

While microgel production using droplet microfluidics, including droplet generation, in-drop polymerization, and transfer of beads from oil to an aqueous solution, is conceptually straightforward, achieving reproducible and predictable control over their physical properties remains challenging. This is largely due to the extreme sensitivity of microscale reactors to small fluctuations in flow rates, reagent concentrations and interfacial transport dynamics. Key parameters, such as flow rates, device geometry and pre-gel composition (monomer [39], crosslinker [18] and initiator [33] concentrations), influence size and elasticity of the microgels. Additional phenomena arising from the interplay of the mentioned parameters may influence polymerization kinetics and network architecture. For instance, an interplay between the pre-gel droplet size and initiator concentration has been found to affect polymerization efficiency, with a critical threshold initiator concentration necessary for efficient in-drop polymerization [40]. However, even when polymerization appears successful, small changes in local reaction conditions can produce microgels with distinct network architectures and mechanical behavior. Moreover, controlling and decoupling the size and elasticity remain challenging as the pre-gel droplets of identical size but different pre-gel composition result in microgels with varying sizes due to elasticity-dependent swelling [39].

In addition to these factors, the microfluidic environment itself introduces additional complexities. Unlike bulk gels, where the catalyst is uniformly distributed, polymerization in microgels occurs upon catalyst diffusion from the continuous phase into pre-gel droplets. This creates interfacial transport phenomena and localized concentration gradients that are highly sensitive to flow dynamics and microscale mass transfer. These often-overlooked factors can substantially influence polymerization kinetics and, ultimately, the physical properties of the resulting microgels. In this study, we address this critical but underexplored aspect by systematically varying flow conditions during droplet production and examining their effect on the physical properties of the resulting microgel beads. To the best of our knowledge, our findings reveal for the first time that microgel elasticity is strongly influenced by flow conditions, even when pre-gel droplet composition is held constant, highlighting the critical role of microscale transport processes in achieving standardized production of cell-mimicking microgels.

A flow-focusing microfluidic chip was employed to generate PAAm pre-gel droplets dispersed in a continuous oil phase, with a fixed composition of both phases. The droplets were subsequently polymerized and transferred to an aqueous phase to yield microgel beads. To isolate the impact of flow conditions without introducing droplet size-related influences on bead polymerization, a unique droplet production regime was identified in which the mean droplet diameter (∼13 µm) varied by less than 1 µm across different flow conditions. This was achieved by keeping total flow rate constant while varying the PAAm-to-oil flow rate ratio. Despite comparable pre-gel droplet sizes and identical pre-gel composition, the resulting beads exhibited unexpected variation in mean Young’s modulus (0.7 to 1.8 kPa). Two distinct production regimes were identified, each affecting bead size and elasticity. We investigated the underlying cause of this behavior and identified the transport of the catalyst from oil to aqueous solution as a key driver for this variation. Our findings demonstrate that flow conditions can indirectly regulate in-drop polymerization by modulating interfacial transport of the catalyst. This highlights a critical yet often neglected factor in microgel production, providing essential insights into the microfluidic parameters required for standardized and reproducible fabrication of PAAm microgels with controlled size and elasticity. Importantly, we identify a novel strategy to fine-tune the microgel elasticity from a fixed pre-gel composition solely through flow control, eliminating variability associated with pipetting or preparing different pre-gel mixtures. This offers a robust and straightforward technique to match microgel elasticity to the mechanical properties of specific cell types. Taken together, our results highlight how sensitively the microfluidic parameters shape PAAm microgel properties and advance efforts towards reproducible, application-ready microgel systems for use in mechanobiology, standardization of novel diagnostic methods, tissue engineering, and drug delivery.

## Results & discussion

### 1. Polyacrylamide pre-gel droplet production

The versatility of the microgel beads has made them indispensable tools for diverse applications, including cell mechanics studies, deformability cytometry and generation of 3D scaffolds [17,25,27]. Achieving precise control over their size and elasticity is paramount to tailoring their properties to meet specific application needs as well as for ensuring reliable and reproducible experimental outcomes. Microgel beads are formed by various techniques such as batch emulsification, lithography and mechanical fragmentation of bulk hydrogels; however, these approaches often lack precise control over final bead size and monodispersity or suffer from low throughput [13,31,41]. In contrast, droplet microfluidics enables unparalleled control over the high-throughput generation of monodisperse droplets containing pre-gel polymer components, whose size and composition directly define the size and elasticity of the resulting microgel beads [37,42]. This process differs fundamentally from bulk polymerization, as the polymerization within the confined microenvironment of the rapidly moving pre-gel droplets can be influenced by interfacial effects, concentration gradients and flow dynamics [23]. Additionally, flow rates during droplet formation can significantly impact droplet size, monodispersity and reactant distribution and mixing, which can further shape the size and elasticity of the final beads [43,44]. To address this, we investigated the influence of flow conditions on the physical properties of the microgel beads. As a first step, we focused on identifying the optimal experimental set-up, microfluidic chip design and flow parameters to stably and reproducibly generate monodisperse pre-gel droplets of the desired size, suitable for a range of applications in mechanobiology.

#### 1.1 Microfluidic set-up for PAAm pre-gel droplet production

To investigate the influence of flow conditions on the physical properties of the microgel beads, we developed a microfluidic platform for the reliable and stable production of monodisperse PAAm pre-gel droplets dispersed in an oil phase. The system utilized a PDMS-based flow-focusing microfluidic chip, interfaced with a pressure-based microfluidic controller (Fluigent MFCS™ EX) equipped with two flow sensors (FS) and a closed-loop feedback system. Controlled via the OxyGEN (Fluigent) software, the system automatically adjusts pressures to maintain constant flow rates during droplet production (Figure 1A, top). This setup enabled precise regulation of flow rates and rapid flow stabilization, minimizing fluctuations compared to conventional syringe-pump systems. Unlike syringe pumps, which often introduce pulsing effects and mechanical drift that compromise droplet size uniformity, the pressure-based system ensured pulse-free flow. As a result, it maintained consistent flow rates throughout production, ensuring uniform droplet size and reproducibility across experiments [45,46]. The pressure controller was connected to a nitrogen gas supply to pressurize the vials with an inert gas, preventing the degradation of free radicals from the initiator caused by oxygen exposure during the droplet production process [47,48].

**Figure 1.**
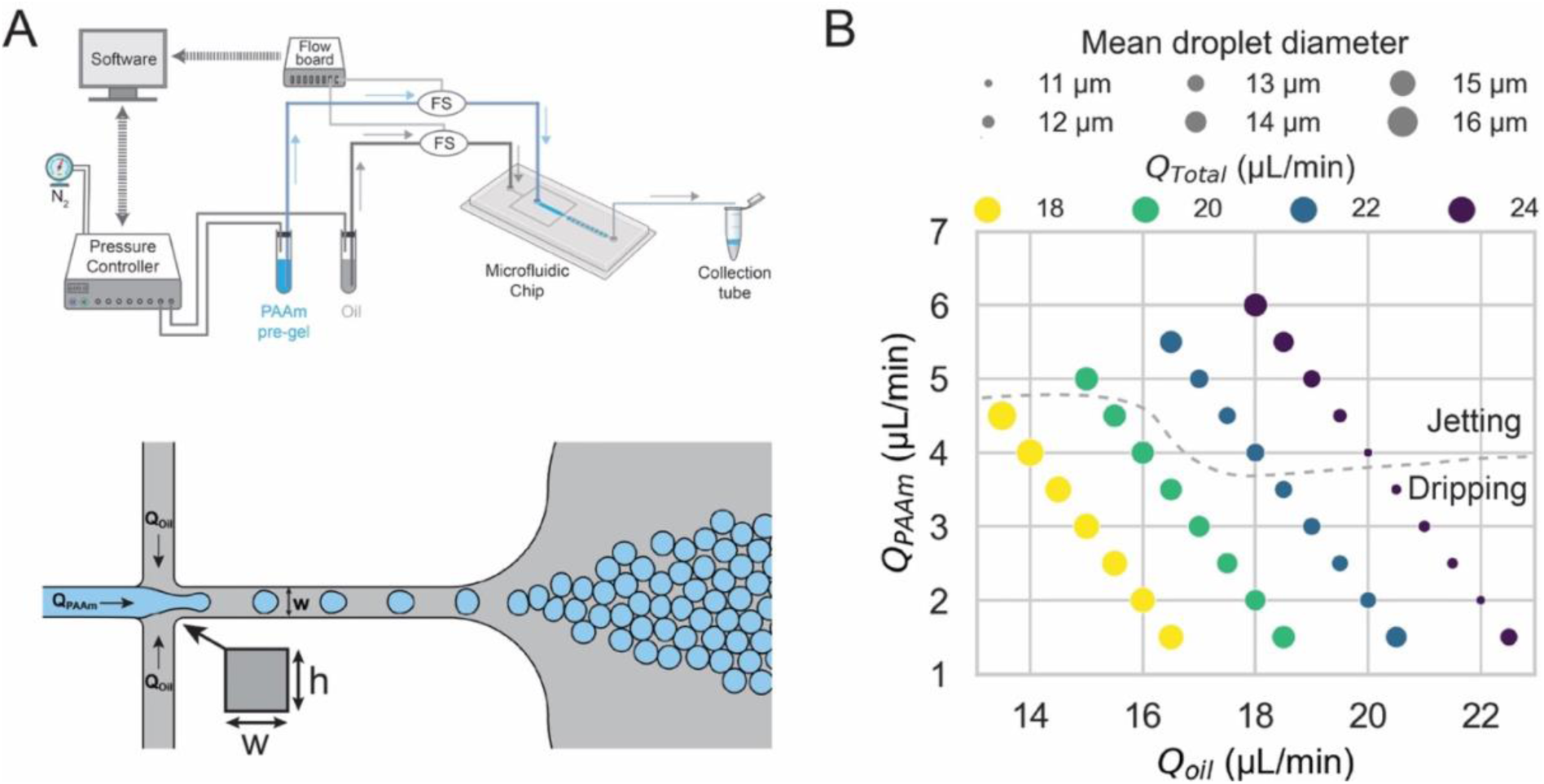
Optimization of polyacrylamide pre-gel droplet production. **(A)** Schematic illustration of the microfluidic platform (top) used for the production of polyacrylamide (PAAm) pre-gel droplets. The setup includes a pressure controller that pressurizes vials containing PAAm pre-gel mixture and oil solutions. These vials are connected to the microfluidic chip via tubing, including two flow sensors (FS), enabling constant flow conditions for both solutions through a closed-loop feedback control system between the pressure controller and the FS. The generated droplets are subsequently collected in the collection tube. The lower panel depicts a schematic representation of PAAm pre-gel droplet production process within the flow-focusing microfluidic chip. Droplet formation is driven by the dispersed phase flow rate (*Q_PAAm_*) and the continuous phase flow rate (*Q_Oil_*). The crossjunction region features a square channel with a width of *w* = 15 µm. **(B)** Variation of pre-gel droplet mean diameter as a function of *Q_PAAm_* and *Q_Oil_* for different values of the total flow rate *Q_Total_* (= *Q_PAAm_* + *Q_Oil_*). Each data point (circle) corresponds to specific values of *Q_PAAm_*, *Q_Oil_* and *Q_Total_*. The size of the circles reflects the mean droplet diameter. The dashed line separates the two droplet production regimes: dripping and jetting.

The dispersed phase consisted of a PAAm pre-gel mixture, including acrylamide (AAm) as a monomer, bis-acrylamide (BIS) as a crosslinker and ammonium persulfate (APS) as a radical initiator. By adjusting the pre-gel composition, the elasticity of the microgel beads can be tuned. A pre-gel mixture with a total monomer concentration [34],

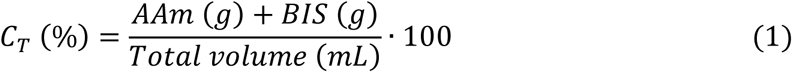

of 5.6% w/v was chosen to achieve beads with elasticity around 1 kPa, which is of interest for several biophysical applications [7,49,50].

The continuous phase contained a solution of fluorinated oil (HFE-7500) with ammonium Krytox® as a surfactant and TEMED as a catalyst. The surfactant stabilized the emulsion, preventing droplet aggregation, while TEMED, upon diffusing into the pre-gel droplets, catalyzed the free-radical polymerization process. In bulk gel systems, the polymerization can proceed in the absence of TEMED, albeit significantly delayed [51], highlighting the critical role of TEMED in facilitating efficient polymerization. In contrast, in microdroplets, polymerization was unsuccessful even after 24 h and at standard polymerization temperature (i.e. 65°C) without TEMED, likely due to rapid degradation of APS in the confined volume and increased oxygen diffusion at the droplet’s oil–water interface, both of which inhibit free-radical formation [30].

Both the PAAm pre-gel and oil solutions were loaded into separate vials and connected to the chip’s inlets via FEP tubing. The microfluidic chip featured a flow-focusing device geometry with a crossjunction region, where both the channel width (*w*) and height (*h*) measured 15 µm (Figure 1A, bottom). This geometry was designed to produce pre-gel droplets that eventually form microgel beads with a diameter of around 15 µm, suitable for various applications in mechanobiology [17,22,25,27,52]. The flow rates of both the PAAm pre-gel mixture (*Q_PAAm_*) and oil solution (*Q_Oil_*) were controlled by pressurizing the corresponding vials. A closed-loop feedback system adjusted the pressure in real time to maintain constant flow rates during droplet production (Figure S1). This compensated for possible variations in channel resistance due to partial filter blockage, localized or incomplete channel clogging by debris, slight leakage at tubing connections, or partial chip delamination. It was essential for ensuring the production of monodisperse droplets and maintaining reproducibility across experiments. Droplet formation occurred at the crossjunction, where the PAAm stream was pinched by two lateral oil streams, generating uniform droplets under controlled flow conditions. These pre-gel droplets were then collected in a tube connected to the chip’s outlet chamber using the same type of FEP tubing (Figure 1A, top).

The diameter of droplets in droplet-based microfluidics is predominantly influenced by parameters such as the flow rates of the dispersed and continuous phases, chip geometry, fluid viscosities and interfacial tension, as shown in numerous liquid-liquid systems [53–58]. Using pre-gel mixture as the dispersed phase introduces additional complexities, as variations in its components alter both the viscosity and the interfacial tension. These changes directly influence droplet formation and stability [57,58]. This was evident in our system, where using a pre-gel solution in buffer versus buffer alone as the dispersed phase, under fixed chip geometry and flow parameters, resulted in different droplet sizes (Figure S2). Specifically, droplets exhibited smaller diameters when pre-gel components were added to the buffer, mostly due to the increased solution viscosity. Independent of the dispersed phase composition, the mean droplet diameter (*D̄*_*Droplet*_) increased more significantly with an increase in the total flow rate (*Q_Total_* = *Q_PAAm_*+ *Q_Oil_*), and showed a slight increase with an increase in the flow rate ratios (*Q_PAAm_*/*Q_Oil_*). This highlights how the pre-gel mixture composition can affect droplet size and underscores the importance of identifying flow parameters that minimize droplet size variation. To eliminate droplet size variation caused by differences in pre-gel composition, we maintained a constant total monomer concentration across all experiments (*C_T_* = 5.6% w/v). A previous study reported that for droplets containing the same pre-gel composition, differences in droplet diameter led to size-dependent discrepancies in polymerization efficiency [40]. To minimize the impact of droplet size on the free-radical polymerization and to reliably assess the influence of flow conditions on in-drop polymerization, we fixed the chip geometry and the composition of both the dispersed and continuous phases. We then investigated flow parameters that enable the consistent production of monodisperse pre-gel droplets with minimal size variation across different flow rate conditions.

#### 1.2 Optimization of monodisperse pre-gel droplet production with minimal size variation

We aimed to identify specific combinations of *Q_PAAm_* and *Q_Oil_* that consistently produce monodisperse pre-gel droplets with minimal diameter variation, despite changes in flow rates. This approach enabled a systematic study of the influence of flow parameters on pre-gel droplet formation and final microgel bead properties, while eliminating potential uncertainties related to droplet size variation. Using a fixed device geometry and constant PAAm pre-gel composition, we varied both the *Q_PAAm_* and *Q_Oil_* to identify the optimal flow rate combinations for producing monodisperse beads (∼15 µm in diameter), suitable for various applications in biophysics. Droplet break-off regimes were recorded at the flow-focusing crossjunction region using a high-speed camera (3000 fps), while the droplet diameter distribution was analyzed by recording videos downstream of the crossjunction region.

We observed two distinct droplet break-off regimes: dripping and jetting, separated by a dashed line in Figure 1B. Consistent with previous observations, in the dripping regime, the PAAm pre-gel stream was pinched off into monodisperse droplets directly at the crossjunction. Whereas in the jetting regime, PAAm stream extended further downstream, resulting in polydisperse pre-gel droplet diameters (Figure S3) [53,59]. This transition between dripping and jetting regimes occurred in two scenarios: (*i)* for a fixed *Q_Oil_*, increasing the *Q_PAAm_* enhanced the inertial forces of the PAAm jet, but the shear forces from the oil phase were insufficient to break the PAAm stream into droplets, leading to jetting, (*ii*) for a fixed *Q_PAAm_* (4 or 4.5 µL/min), increasing *Q_Oil_* introduced stronger viscous drag forces on PAAm stream, transitioning from dripping to jetting regime [60] (Figure 1B).

For a fixed *Q_PAAm_*, increasing *Q_Oil_* led to a decrease in *D̄*_*Droplet*_ due to increasing drag forces from the oil solution, which accelerated the droplet break-up [39,54], (Figure 1B and S3). Consistent with previous findings, increasing the *Q_Total_* from 18 to 24 µL/min resulted in a decrease in *D̄*_*Droplet*_ from 16.2 µm to 11.6 µm [61] (Figure 1B). Notably, for a fixed *Q_Total_* ≤ 22 µL/min, the *D̄*_*Droplet*_ showed minimal variation across varying flow rate ratios (*Q_PAAm_*/*Q_Oil_*). For instance, with fixed *Q_Total_* values of 18, 20 and 22 µL/min, *D̄*_*Droplet*_ varied by only 8%, 6% and 11% respectively, despite changes in *Q_PAAm_*/*Q_Oil_* ratios (Figure 1B). A similar trend was observed by Dewandre et al. [62], who reported that varying the dispersed phase flow rate, while keeping the continuous phase flow rate constant in a co-flow-focusing geometry, led to a maximum droplet size variation of about 15%, along with an increase in droplet production frequency. The small droplet size variation was attributed to the dominance of the viscous forces from the continuous phase, suggesting that increasing the dispersed phase flow rate primarily influenced the production frequency rather than droplet diameter. Here, we simultaneously varied both the *Q_PAAm_* and *Q_Oil_* across different ratios, introducing subtle changes in inertial forces from *Q_PAAm_* and viscous forces from *Q_Oil_*. These small shifts led to only minor variations in *D̄*_*Droplet*_ under fixed *Q_Total_*. Pre-gel droplets produced at *Q_Total_* = 20 µL/min consistently exhibited the desired size range (13–14 µm) with minimal size variation across most *Q_PAAm_*/*Q_Oil_* ratios and remained stable within the dripping regime. Therefore, this flow condition was selected for detailed investigation of droplet diameter and production frequency.

#### 1.3 Influence of *Q_PAAm_*/*Q_Oil_* ratio on droplet production

The *Q_PAAm_*/*Q_Oil_* ratio was varied from 0.08–0.38, keeping the chip geometry, PAAm pre-gel composition (*C_T_* = 5.6%) and *Q_Total_* (= 20 µL/min) constant. The droplet diameter (*D*_*Droplet*_) was determined by analyzing frames extracted from videos recorded during droplet production. The droplet production frequency (*f*) was calculated as,

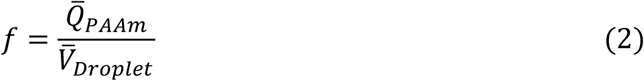

where, *Q̄*_*PAAm*_ is the mean flow rate of PAAm pre-gel solution recorded by the flow sensor during production, and *V̄*_*Droplet*_ is the mean droplet volume.

Within the dripping regime (*Q_PAAm_*/*Q_Oil_* = 0.08–0.29), different behaviors in *D̄*_*Droplet*_ were observed (Figure 2A). Specifically, when *Q_PAAm_*/*Q_Oil_* increased from 0.08 to 0.21, the *D̄*_*Droplet*_ increased from 13.0 ± 0.2 µm to 13.8 ± 0.2 µm, representing a maximum variation of 6%. A further increase in *Q_PAAm_*/*Q_Oil_* from 0.21 to 0.29 resulted in a slight variation of 0.7% in *D̄*_*Droplet*_ (to 13.9 ± 0.2 µm). Droplets formed in the dripping regime exhibited a narrow size distribution, with a coefficient of variation (*C.V.*, defined as the ratio of the standard deviation to *D̄*_*Droplet*_) below 1.5%, confirming highly monodisperse droplet generation [63]. As previously discussed, the observed small increment in *D̄*_*Droplet*_ is attributed to a simultaneous increase in inertia of *Q_PAAm_* and reduction in viscous shear forces from *Q_Oil_* with increasing *Q_PAAm_*/*Q_Oil_* ratio [60,64]. In contrast, this interplay between inertia and shear forces showed a pronounced effect on *f*, increasing from approximately 22 kHz to 54 kHz, nearly 2.5–fold increase with increasing *Q_PAAm_*/*Q_Oil_* ratio in the dripping regime (Figure 2).

**Figure 2.**
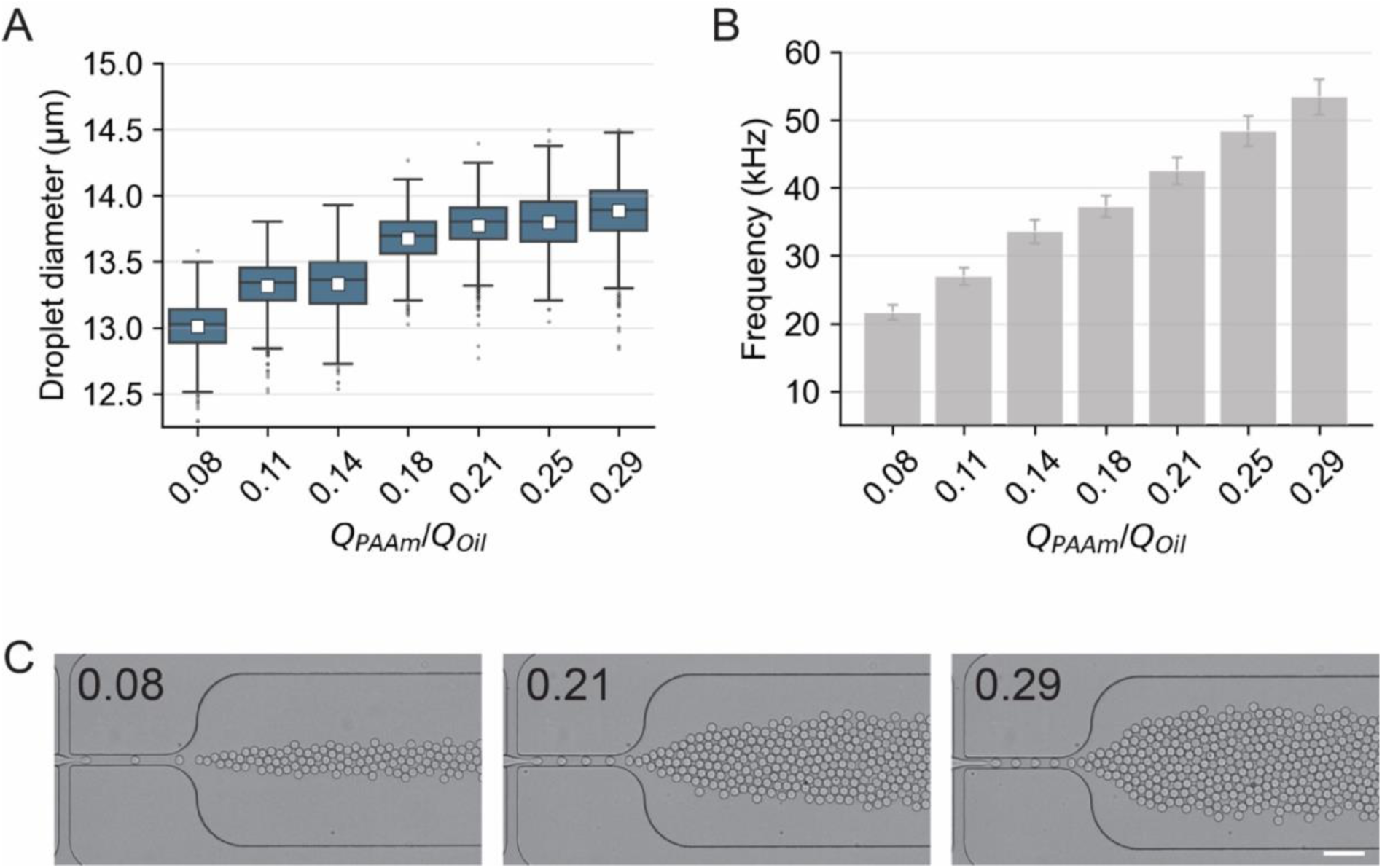
Influence of *Q_PAAm_/Q_Oil_* ratio on pre-gel droplet production. Droplet diameter and production frequency as a function of *Q_PAAm_/Q_Oil_* ratios for a fixed *Q_Total_*= 20 µL/min. **(A)** The box plots represent the droplet diameter distribution at varying *Q*_*PAAm*_⁄*Q*_*Oil*_. Each box represents the interquartile range (IQR) covering data between 25^th^ and 75^th^ percentiles. The horizontal line within the box represents median, while the white square markers indicate mean. Whiskers extend to 1.5 times the IQR, while the data points outside this range are represented as dots, which indicate outliers. **(B)** The bar plot depicts the droplet production frequency (kHz) as a function of *Q_PAAm_/Q_Oil_*. The frequency was calculated as the ratio of *Q̄*_*PAAm*_ and mean droplet volume, *V̄*_*Droplet*_, while the error bars represent the propagated uncertainty based on the standard deviation of the *Q̄*_*PAAm*_ and *V̄*_*Droplet*_. **(C)** Bright-field images showing pre-gel droplet formation at the crossjunction for *Q_PAAm_/Q_Oil_* values of 0.08, 0.21 and 0.29 (indicated in the top-left corner of each image), corresponding to the data presented in (A) and (B). Scale bar: 50 µm.

A further increase in the *Q_PAAm_*/*Q_Oil_*(> 0.29) led to droplet break-off transition into jetting regime (Figure S4). In this regime, the reduction in the viscous forces from the oil phase caused PAAm jet radius to widen, leading to uncontrolled break-up due to Rayleigh capillary instability and extended droplet pinch-off time, resulting in less uniform droplet formation and a larger *D̄*_*Droplet*_ [60,64–66]. As a result, we observed an increase in *D̄*_*Droplet*_, with values of 14.5 ± 0.4 µm for *Q_PAAm_*/*Q_Oil_* of 0.33 and 15.2 ± 0.9 µm for *Q_PAAm_*/*Q_Oil_* of 0.38 with *C.V.* of up to 5.9%. Meanwhile, the *f* did not vary significantly, with values of 53 kHz and 50 kHz for the *Q_PAAm_*/*Q_Oil_* ratios of 0.33 and 0.38, respectively (Figure S4).

Overall, our results showed that, for a fixed chip geometry, pre-gel composition and *Q_Total_*, a rational increase of the *Q_PAAm_*/*Q_Oil_* ratio influenced the droplet break-off regimes, with minor variations in droplet size. Under the dripping regime, *D̄*_*Droplet*_ varied by less than 1 µm, maintaining high monodispersity (C.V. < 1.5%), while *f* increased linearly up to 54 kHz. The pre-gel droplets produced under these flow conditions were collected, polymerized and transferred to a Phosphate-Buffered Saline solution (1×PBS) after multiple washing steps. The polymerized microgel beads in 1×PBS will be larger than in oil, as the hydrogel meshwork tend to swell upon absorbing water.

### 2. Influence of *Q_PAAm_*/*Q_Oil_* ratio on pre-gel droplet polymerization and physical properties of resulting PAAm beads

Unlike bulk hydrogels, microgel fabrication introduces size as a critical parameter, which is determined not only by the templating droplet diameter but also by the pre-gel composition and polymerization conditions. Upon transfer into aqueous solution, microgels swell due to water uptake into the polymer meshwork, a process governed by crosslinking density. Higher crosslinking limits swelling, leading to smaller beads, while lower crosslinking allows greater swelling and results in larger beads [67–69]. Even with identical droplet diameters, variations in monomer [39], crosslinker [18,23,32], or initiator [33] concentrations influence swelling behavior and ultimately determine the final bead size. Given the identical pre-gel composition of the droplets used in our study, the microgel beads were expected to follow a size trend similar to the corresponding pre-gel droplets and to exhibit comparable meshwork formation affecting their swelling behavior and mechanical properties. To verify this, we characterized the physical properties of the microgels polymerized from the pre-gel droplets shown in Figure 2 and S4, by analyzing their polymerization efficiency, diameter, swelling, dry mass and elasticity.

#### 2.1. Polymerization efficiency of the PAAm pre-gel droplets

To evaluate the polymerization efficiency of the pre-gel droplets, the microgel bead production yield (%) was calculated as follows,

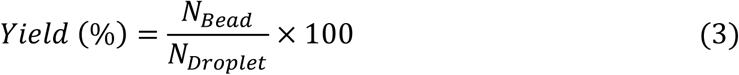

where *N*_*Bead*_ is the measured number of beads polymerized and *N*_*Droplet*_ is the estimated number of droplets produced. *N*_*Droplet*_ was calculated as the ratio of the volume of PAAm pre-gel solution used to produce droplets (*V*_*PAAm*_= 50 µL) to the mean volume of droplets (*V̄*_*Droplet*_). Since the droplet production frequency varied with the *Q_PAAm_*/*Q_Oil_* ratio, we fixed *V*_*PAAm*_ at 50 µL for each tested flow condition, rather than fixing the collection time, to maintain a relatively consistent *N*_*Droplet*_ across different production conditions (Figure S5). The yield increased with rising *Q_PAAm_*/*Q_Oil_* ratios, showing the largest variation (from 0.2% to 44.2%) within the dripping regime and then nearly stabilized in the jetting regime, with values of 42% and 50% at *Q_PAAm_*/*Q_Oil_* ratios of 0.33 and 0.38, respectively (Figure 3A).

**Figure 3.**
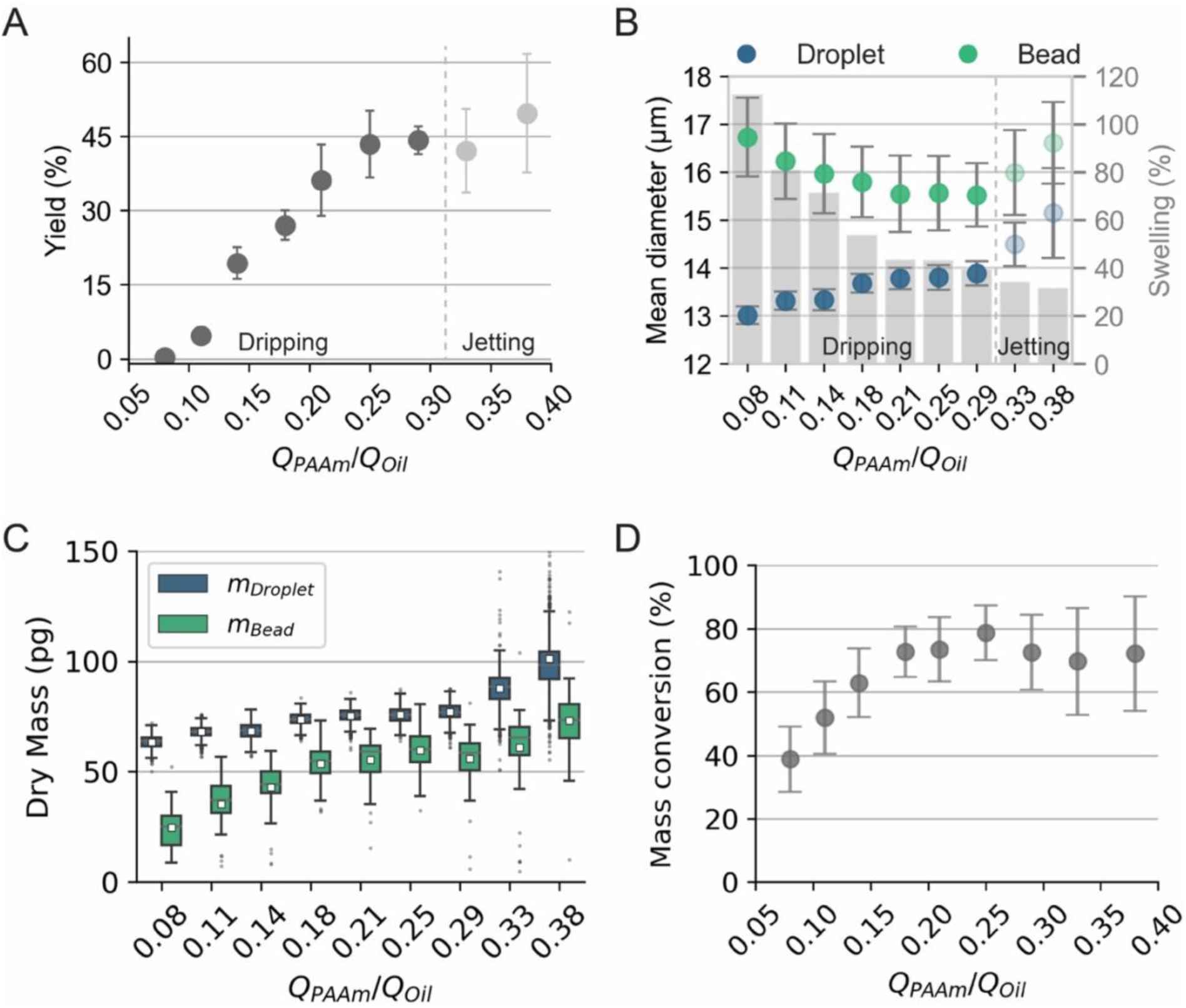
Influence of *Q_PAAm_/Q_Oil_* on pre-gel droplet polymerization and physical properties of resulting beads. Analysis of the in-drop polymerization efficiency and the physical properties of microgel beads produced under constant *Q_Total_* = 20 µL/min and varying *Q_PAAm_/Q_Oil_*. **(A)** Polymerization yield (%) as a function of *Q_PAAm_/Q_Oil_*. The yield (%) was calculated as the ratio of the measured number of polymerized beads (*N*_*Bead*_) to the estimated number of droplets produced (*N*_*Droplet*_). Data points represent mean values and the error bars represent the propagated uncertainty derived from standard deviation of *N*_*Droplet*_ and *N*_*Bead*_. The dashed line separates the two droplet production regimes: dripping and jetting. **(B)** Comparison of mean droplet diameter (blue) and mean bead diameter (green) for varying *Q_PAAm_/Q_Oil_* ratios. The error bars represent the standard deviation. The bar plot shows the mean bead swelling (%) for each *Q_PAAm_/Q_Oil_* ratio, calculated as the volume increase relative to the droplet volume. The dashed line separates the two droplet production regimes: dripping and jetting. **(C)** Comparison between the estimated dry mass of pre-gel components in droplets (*m*_*Droplet*_, in blue) and dry mass of the polymer in the beads (*m*_*Bead*_, in green) measured using Optical Diffraction Tomography as a function of *Q_PAAm_/Q_Oil_*. Each box represents the interquartile range (IQR) covering data between 25^th^ and 75^th^ percentiles. The horizontal line within the box represents median, while the white square markers indicate mean. Whiskers extend to 1.5 times the IQR, while the data points outside this range are represented as dots, which indicate outliers. **(D)** Mass conversion (%) as a function of *Q_PAAm_/Q_Oil_*. Data points and error bars represent mean and S.D., respectively.

These results suggest a strong link between flow conditions and polymerization efficiency of the pre-gel droplets. DiLorenzo et al. reported size-dependent discrepancies in the ability of microgel formation from microdroplets of identical pre-gel composition, where reduction in droplet diameter by ∼50% led to failed polymerization [40]. Specifically, they demonstrated that the pre-gel droplets produced with *C_T_* ∼6.3 % and APS concentration of 4 g/L polymerized effectively at a diameter of about 50 µm, while the droplets ∼25 µm in diameter failed to polymerize due to the necessity of higher APS concentration (i.e. 5 g/L). This was attributed to the critical balance between heat generation and dissipation during the exothermic polymerization reaction. In our study, the pre-gel droplets were produced at a fixed *C_T_* = 5.6%, APS concentration of 8.25 g/L, and *D̄*_*Droplet*_ ranging between 13–14 µm (Figure 2A). These values lie well beyond the minimum APS concentration necessary (i.e. ∼7 g/L) for polymerization success. Based on this, we expected the same polymerization efficiency for the microgel beads produced in our study under different flow conditions. However, we observed a ∼220-fold increase in yield as the *Q_PAAm_*/*Q_Oil_* increased, despite maintaining the APS concentration above the critical threshold within pre-gel droplets comparable in size. The observed differences in polymerization efficiency suggest variations in the polymerization process, which may have influenced the meshwork formation. This, in turn, could affect the beads’ water uptake capacity and ultimately impact their physical properties, such as diameter and swelling behavior.

#### 2.2. Bead diameter and swelling behavior as a function of *Q_PAAm_*/*Q_Oil_*

The diameter of the beads (*D*_*Bead*_) suspended in 1×PBS was measured by analyzing brightfield microscopy images of the beads using Fiji. The mean bead radius (*r̄*_*Bead*_) and mean radius of the corresponding pre-gel droplets (*r̄*_*Droplet*_) were used to calculate the swelling degree of the beads as follows [70]:

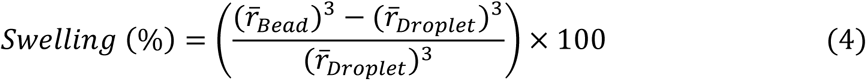

Since all the pre-gel droplets were generated from the same composition and polymerized under identical conditions, we initially expected the resulting microgel beads to exhibit consistent swelling behavior and bead diameters to vary in line with those of the pre-gel droplets. Contrary to this expectation, we observed a clear dependence of bead swelling on the *Q_PAAm_*/*Q_Oil_*, resulting in an unexpected trend in bead diameter variation. In the dripping regime (*Q_PAAm_*/*Q_Oil_* = 0.08–0.29), the resulting beads exhibited two distinct behaviors in diameter variation (Figure 3B). As the *Q_PAAm_*/*Q_Oil_* increased from 0.08 to 0.21, the mean bead diameter (*D̄*_*Bead*_) decreased from 16.7 ± 0.8 µm to 15.5 ± 0.7 µm, showing an opposite trend compared to the size variation of the corresponding pre-gel droplets. This counterintuitive trend can be attributed to differences in key structural properties of the formed polymer meshwork, such as the degree of crosslinking, network architecture, and its elasticity, affecting solvent imbibition [68,69,71]. This was corroborated by the measured swelling degree, which decreased from 112% to 44%. The reduced swelling at higher *Q_PAAm_*/*Q_Oil_* (0.21) suggests an increased crosslinking density and a more tightly formed polymer network, leading to a smaller bead size. For *Q_PAAm_*/*Q_Oil_* = 0.21–0.29, *D̄*_*Bead*_ remained constant at about 15.5 µm, indicating minimal variation in the meshwork structure. In this flow ratio range, swelling decreased slightly from 43% to 40%, compensating for the slight increase in pre-gel droplet size and yielding beads with comparable size. In the jetting regime, the swelling decreased further to 34% and 32%, respectively, for *Q_PAAm_*/*Q_Oil_* of 0.33 and 0.38, suggesting the formation of an even more highly crosslinked polymer network. In this regime, bead diameters followed the trend of the corresponding pre-gel droplet sizes, with *D̄*_*Bead*_ of 16.0 ± 0.9 µm and 16.6 ± 0.9 µm for *Q_PAAm_*/*Q_Oil_* of 0.33 and 0.38, respectively. As expected, beads produced in the dripping regime were more monodisperse (C.V. < 5%) compared to those produced under the jetting regime (C.V. > 5%).

#### 2.3. Polymer network formation efficiency as a function of *Q_PAAm_*/*Q_Oil_*

The unexpected trends observed in the bead diameters and swelling behavior indicated the differences in the underlying polymer architecture and crosslinking density, despite identical pre-gel composition of the droplets produced across *Q_PAAm_*/*Q_Oil_* ratios. Although the initial concentrations of monomer and crosslinker are precisely defined, the extent to which these pre-gel components are incorporated into the microgel network remains unclear. This, in turn, determines the final physical properties of the microgels. To better understand these differences, measuring the dry mass of individual microgel beads and comparing it to the mass of AAm and BIS present in the original pre-gel droplets can provide valuable mechanistic insights into the swelling trends and size variations observed across *Q_PAAm_*/*Q_Oil_* ratios. Here, we utilized optical diffraction tomography (ODT), which acquires 3D tomograms of the refractive index (RI) distribution within single beads, enabling precise calculation of their dry mass and mass density [72]. The dry mass values of the beads measured by ODT (*m*_*Bead*_) were compared to the estimated dry mass of the corresponding pre-gel droplets (*m*_*Droplet*_, calculated as *m*_*droplet*_ = *C*_*T*_ × *V*_*Droplet*_, assuming a constant *C*_*T*_ in each droplet) (Figure 3C).

The analysis of the dry mass measurements revealed three key trends: *i*) The mean dry mass of the beads (*m̄*_*Bead*_) was consistently lower than the mean dry mass of the pre-gel droplets (*m̄*_*Droplet*_). This difference can be attributed to incomplete monomer conversion during free-radical polymerization, influenced by termination reactions, initiator decomposition [73–75] and microgel-specific factors such as oxygen inhibition at the droplet interface and rapid reaction kinetics in small volumes [76]. The variations in *m̄*_*Droplet*_ were directly correlated to the variations in the droplet volume as reported in Figure 2A and S4. (*ii*) The *m̄*_*Bead*_ also showed dependence on the *Q_PAAm_*/*Q_Oil_* ratios. It increased by 120%, from ∼25 pg to ∼55 pg as the *Q_PAAm_*/*Q_Oil_* increased from 0.08 to 0.21, while it increased only by 9% to a maximum of ∼60 pg as *Q_PAAm_*/*Q_Oil_* increased to 0.29. Beads produced at *Q_PAAm_*/*Q_Oil_* = 0.33 and 0.38 exhibited *m̄*_*Bead*_ of ∼61 pg and ∼73 pg, respectively. *iii*) While the trend in *m̄*_*Bead*_ aligned with the increasing *m̄*_*Droplet*_, the magnitude of increment was significantly different across *Q_PAAm_*/*Q_Oil_* ratios. This suggested that the initial monomers were converted into a stable polymer network with varying efficiency, which can be quantified using the mass conversion, defined as follows:

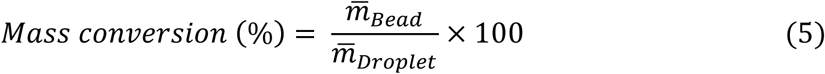

The analysis revealed a linear increase in mass conversion from ∼39% to ∼73% with the increase in *Q_PAAm_*/*Q_Oil_* from 0.08 to 0.18 (Figure 3D). For higher values of the *Q_PAAm_*/*Q_Oil_*, it remained constant, with mean values varying between approximately 70% and 80%. The maximum mass conversion of ∼80% was obtained for *Q_PAAm_*/*Q_Oil_* = 0.25. Previous studies on bulk PAAm hydrogels have demonstrated monomer conversion exceeding 95% [34,73,74,77]. The lower mass conversion in pre-gel droplets can be ascribed to the unique polymerization conditions in microdroplets [78,79], where the high surface-to-volume ratio favors fast oxygen diffusion, promoting radical scavenging [40,48] and fast heat dissipation that can contribute to the formation of an inhomogeneous meshwork. The measurement of bead dry mass and volume using ODT also enabled the calculation of the final bead density (Figure S6), a relevant parameter in several microfluidic applications, such as flow-based separation, focusing or sedimentation [80,81]. From the tomograms presented in Figure S7, no significant variation in RI along the bead radius was observed, suggesting that the crosslinking density was uniform across the bead diameter. This ruled out a core–shell structure in the beads produced under different flow conditions, which could have potentially arisen due to TEMED diffusion. While further analysis of the RI standard deviation within each bead may reflect fine structures and irregularities within the meshwork of individual beads, the current spatial resolution limits of the ODT system (121 nm lateral and 444 nm axial) restricts definitive conclusions. Nonetheless, analyzing the mechanical properties of the microgels can provide further insights into the structural characteristics of the beads produced under varying flow conditions.

#### 2.4 Influence of *Q_PAAm_*/*Q_Oil_* on the mechanical properties of the PAAm microgel beads

The combined analysis of the microgel bead size, swelling and dry mass revealed a clear dependence of their structural properties on the *Q_PAAm_*/*Q_Oil_* used during pre-gel droplet production. Specifically, beads obtained from droplets produced at lower *Q_PAAm_*/*Q_Oil_* exhibited larger sizes, higher swelling and lower dry mass compared to those generated at higher ratios. Based on these observations, a variation in the mechanical properties of the network is expected as a function of the *Q_PAAm_*/*Q_Oil_*, since, according to Flory–Rehner theory [68] and rubber elasticity theory [82], swelling and elastic modulus are intrinsically correlated.

To investigate their mechanical properties, we measured the Young’s modulus (YM) of PAAm microgel beads using real-time deformability cytometry (RT-DC) [83]. Beads were deformed by hydrodynamic shear forces as they flowed through a 30 µm square channel (Figure 4A). Images of the deformed beads were acquired at 3000 fps and analyzed in real-time to detect their contours, which were used to measure both their size and deformation. These values were used to calculate the apparent Young’s modulus for each bead using a lookup table derived from a well-established computational model [84]. A flow rate of 0.12 µL/s, was selected, enabling bead deformation ranging from 0.01 to 0.05, suitable to compute their YM. The deformation analysis of the beads produced in the dripping regime for various *Q_PAAm_*/*Q_Oil_* ratios showed similar area (160–180 µm^2^) and decreasing deformation as the *Q_PAAm_*/*Q_Oil_* increased (Figure 4B). The beads produced under jetting regime showed larger size distribution and area (165–205 µm^2^), with similar deformation independent of the *Q_PAAm_*/*Q_Oil_*. Notably, similar deformation was observed for the beads produced at *Q_PAAm_*/*Q_Oil_* between 0.21 and 0.38.

**Figure 4.**
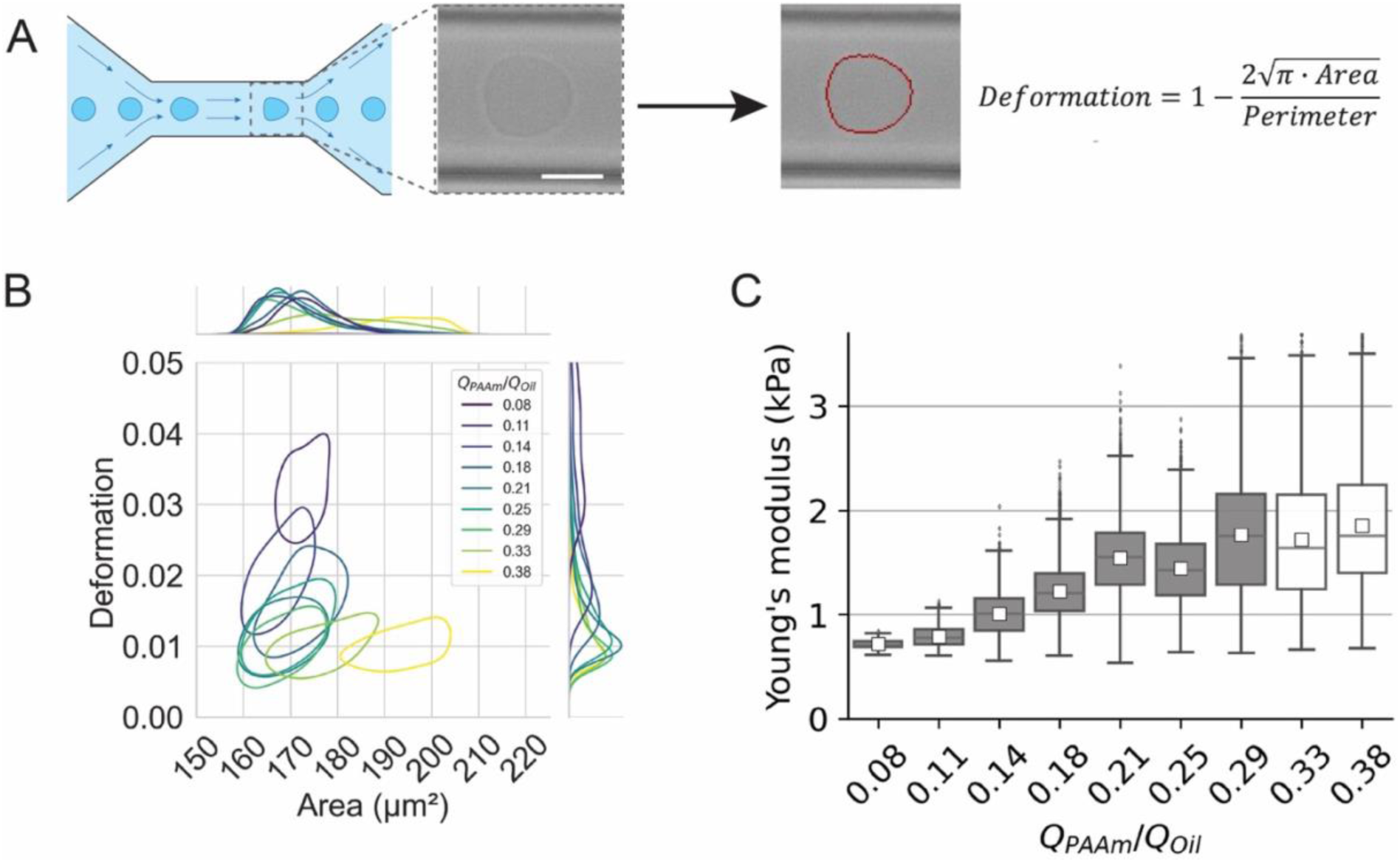
Influence of *Q_PAAm_/Q_Oil_* on the mechanical properties of the PAAm microgel beads. Analysis of the mechanical properties of the microgel beads by real-time deformability cytometry (RT-DC). The beads were obtained by polymerizing the pre-gel droplets produced at a constant *Q_Total_* = 20 µL/min with varying *Q_PAAm_/Q_Oil_*, as presented in Figure 2A and S4. **(A)** Schematic illustration of a bead deformation trajectory while flowing through a 30 µm square microfluidic channel during RT-DC measurement. A bright-field image showing a deformed PAAm bead (left) and a corresponding red contour (right) used for deformation analysis. Scale bar is 10 µm. **(B)** RT-DC analysis of the microgel beads produced at varying *Q_PAAm_/Q_Oil_*, showing contour plots (50% density) of bead deformation vs area alongside corresponding histograms of their deformation and area. **(C)** Box plot showing Young’s modulus distribution of the PAAm microgel beads produced under varying *Q_PAAm_/Q_Oil_*. Each box represents the interquartile range (IQR) covering data between 25^th^ and 75^th^ percentiles. The horizontal line within the box represents median, while the white square markers indicate mean. Whiskers extend to 1.5 times the IQR, while the data points outside this range are represented as dots, which indicate outliers. The color of the boxes shows the two different production regimes: dripping (gray) and jetting (white).

Deformation alone does not fully characterize the mechanical behavior of the beads. The calculation of their YM is needed to account for their ability to deform in a given channel, as larger beads encounter higher hydrodynamic forces compared to smaller ones. The analysis revealed a strong dependence of the YM on the *Q_PAAm_*/*Q_Oil_* (Figure 4C). Specifically, for beads produced at *Q_PAAm_*/*Q_Oil_* ranging from 0.08–0.21, the mean YM increased by 133%, from 0.7 ± 0.1 kPa to 1.5 ± 0.4 kPa. Increasing the *Q_PAAm_*/*Q_Oil_* to 0.25 did not affect bead elasticity. Further increasing the *Q_PAAm_*/*Q_Oil_* led to beads with mean YM of about 1.8 kPa and affected YM distribution, showing higher dispersion of the bead mechanical properties. This trend was in line with the previously observed swelling behavior, showing an inverse relationship between YM and swelling (Figure S8) as reported by Flory-Rehner theory [68,85]. Our analysis showed that the largest variation in YM occurred between *Q_PAAm_*/*Q_Oil_* of 0.08 and 0.21, consistent with the variations observed for bead size, swelling, dry mass, and mass conversion. These results highlighted that, within this flow regime, beads with higher density, obtained for higher *Q_PAAm_*/*Q_Oil_* ratio, exhibited higher Young’s modulus, indicating that the monomers in the corresponding pre-gel droplets were effectively converted into more active elastic chains. *Q_PAAm_*/*Q_Oil_* > 0.21 did not strongly affect the mean values of bead size and elasticity, but primarily increased the standard deviation, indicating greater polydispersity in the physical properties of the beads produced at higher flow ratios.

Taken together, our results showed that with increasing *Q_PAAm_*/*Q_Oil_*, the bead polymerization yield improved, bead swelling decreased and the dry mass in the beads increased (Figure 3). Further mechanical characterization aligned with these trends, showing that the beads produced at higher ratios were consistently stiffer (Figure 4). Collectively, these results, underpinned by fundamental polymer physics, suggest that the interplay between the physical properties of the beads and the fabrication conditions, especially the flow rate ratios, is intrinsically linked to variations in polymerization dynamics of the produced microdroplets. Yet, a perplexing mystery still lingers: how do flow conditions influence the polymerization behavior of the pre-gel droplets so profoundly? What critical, yet elusive factor is responsible for these variations, even when all other conditions are seemingly constant?

### 3. Modulation of interfacial transport by flow rate ratios and its impact on bead polymerization and physical properties

Free-radical polymerization of PAAm bulk gels and their final physical properties have been extensively studied. It is well known that several factors influence gelation kinetics and network formation, including total monomer concentration [34], cross-linker concentration [71,86], temperature [87], catalyst and initiator concentrations [77,88,89] and polymerization methods. Additional factors must be considered when transitioning from bulk hydrogels to microgels fabricated via microfluidic techniques, where interfacial transport phenomena at the microscale can significantly affect polymerization kinetics. Our findings demonstrated that microgel beads formed from the polymerization of pre-gel droplets with identical pre-gel composition under different flow conditions exhibit substantial variations in structural properties. This suggests that polymerization within microfluidic droplets differs from that in bulk gels. Further investigation is needed to identify the key parameters responsible for these variations linked to changes in the *Q_PAAm_/Q_Oil_*.

During the polymerization of microdroplets, interfacial effects play a key role in governing energy and mass transport across both the oil–water and the solid–liquid interfaces [90]. The balance between the heat generated by free-radical polymerization and its dissipation through the oil–water interface affects both the polymerization efficiency and the the resulting polymer meshwork [40]. Similarly, the competition between the diffusion of oxygen and the catalyst (TEMED) into the droplets influences microgel polymerization: oxygen scavenges free radicals and inhibits polymerization [48], while TEMED catalyzes the reaction. All of these interfacial processes are strongly influenced by the droplet size. The heat generated during polymerization scales with droplet volume, while heat dissipation scales with droplet surface area. As a result, larger droplets dissipate heat more slowly (i.e. retain heat longer) than smaller ones, potentially accelerating the reaction rate and improving polymerization efficiency [40]. To assess whether small variations in droplet size were responsible for the observed structural differences, we produced pre-gel droplets with *D̄*_*Droplet*_ ranging from 13 to 15 µm, using two different *Q_PAAm_*/*Q_Oil_* of 0.08 and 0.29. Our analysis showed that both swelling and yield (Figure S9) did not vary with droplet size and were comparable to the values reported in Figure 3. These results showed that diameter variations of less than 1 µm, as tested in our study, are not responsible for the observed structural differences.

Furthermore, the concentration of APS, relative to droplet size, has been shown to impact microgel polymerization. Depending on the droplet size, a critical APS concentration is required for polymerization success. In our study, we used an APS concentration of approximately 8.25 g/L, which is above the critical threshold of 7 g/L reported for droplets of ∼13 µm in diameter [40]. To prevent APS degradation, the pre-gel mixture was pressurized with nitrogen. However, oxygen can still diffuse into the pre-gel mixture and droplets through highly permeable PDMS [91,92] and fluorinated oil [93,94], respectively, which may inhibit microgel polymerization. Prior to droplet formation, oxygen diffusion into the pre-gel mixture can occur at the PDMS-liquid interface depending on the flow rate-dependent residence time of the solution. The volume of pre-gel mixture present in the chip before the cross-junction is approximately 3 µL, resulting in a residence time ranging from 0.5 to 2 min for the highest (5.5 µL/min) and the lowest (1.5 µL/min) *Q_PAAm_* values, respectively. Since this duration is significantly longer than the typical microscale oxygen diffusion timescale (< 1 s), oxygen is expected to affect gel formation regardless of the *Q_PAAm_*/*Q_Oil_*. On the other hand, oxygen permeation continues after droplet formation at the oil–pre-gel interface. This effect becomes more pronounced at lower *Q_PAAm_*/*Q_Oil_*, where fewer droplets are surrounded by a larger oil volume (Figure 2B and S4), providing a possible explanation of the reduced yield observed at low *Q_PAAm_*/*Q_Oil_*(Figure 3A).

Another phenomenon influenced by the flow rate ratio is the diffusion of TEMED from the oil phase into the pre-gel droplets. Since *Q_PAAm_*/*Q_Oil_* directly determines the volume ratio between the total PAAm solution encapsulated in droplets and the surrounding oil phase (i.e., *Q_PAAm_*/*Q_Oil_*= *V_PAAm_*/*V_Oil_*), it also controls the amount of TEMED available per droplet. We observed that lower *Q_PAAm_*/*Q_Oil_* ratios, corresponding to fewer droplets within a larger oil volume, resulted in reduced yield and a lower Young’s modulus of the microgels. This aligns with previous studies showing that higher TEMED concentrations enhance swelling and reduce the bulk modulus of PAAm bulk gels [89]. In our case, the initial TEMED concentration in oil phase (*c*_*T*,*i*_) was constant, but the resulting concentration in the droplets post-diffusion remains unknown. An estimation of the amount of TEMED in each droplet can be obtained by considering the partitioning of TEMED between PAAm and oil phase. The partition coefficient *P* is defined as follows [95]:

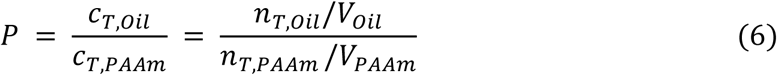

where *c*_*T*,*PAAm*_ and *c*_*T*,*Oil*_ are the TEMED concentration, while the *n*_*T*,*PAAm*_ and *n*_*T*,*Oil*_ are the moles of TEMED in PAAm and oil solutions, respectively, at equilibrium post-production. Due to mass conservation [95,96],

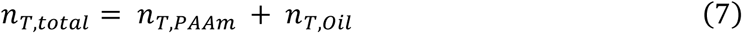

where *n*_*T*,*total*_ is the total number of moles of TEMED initially present only in the oil phase. From equations (6) and (7), moles of TEMED per droplet can be derived as:

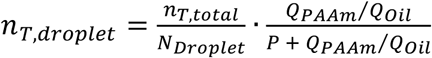

and the concentration of TEMED in a droplet can be calculated as,

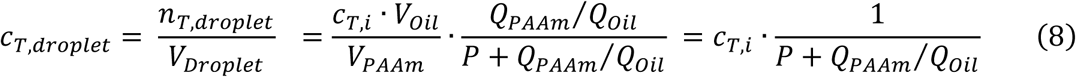

The equation (8) shows how the TEMED concentration per droplet (*c*_*T*,*droplet*_) varies with the *Q_PAAm_*/*Q_Oil_* and the partition coefficient (*P*). TEMED is a polar, water-miscible amine that dissolves better in polar solvents. Given its high solubility in water (>1000 g/L) compared to fluorinated oil (HFE-7500), *P* is expected to be significantly less than 1. A *logP* value of approximately −0.13 has been reported for the n-octanol/water system [97]. Since n-octanol (dielectric constant ∼10.3) is considerably more polar than HFE-7500 (dielectric constant ∼5.8), the *logP* value for the HFE-7500/water system can reasonably be assumed to be much lower than −0.13.

Based on equation (8), we calculated how *c*_*T*,*droplet*_ varied with respect to *Q_PAAm_*/*Q_Oil_* as a function of different *logP* values (Figure 5A). Our analysis showed that *c*_*T*,*droplet*_ became increasingly sensitive to variations in the *Q_PAAm_*/*Q_Oil_* as the partition coefficient decreased. The most pronounced concentration changes occur for *logP* < −1 and *Q_PAAm_*/*Q_Oil_* ≤ 0.21, coinciding with the range in which the physical properties of the microgel beads, including size, swelling, mass and Young’s modulus, exhibited the greatest variation. These results suggested that differences in the amount of TEMED available per droplet can significantly influence the final properties of the microgels. To confirm this, we compared the swelling behavior of beads produced at three different *Q_PAAm_*/*Q_Oil_* values under two experimental conditions, using an oil solution with a fixed TEMED concentration (*c*_*T*,*i*_): *i) Variable V_Oil_ in the collection tube*. As in previously reported conditions (Figure 2A), the amount of TEMED in the collection tube was determined by the *V_Oil_* consumed during production, which varied depending on the *Q_PAAm_*/*Q_Oil_*(see vial pictures in Figure 5A). *ii) Constant V_Oil_ in the collection tube*. During production, the *V_Oil_* in the collection tubes was adjusted across all conditions to match the values obtained at *Q_PAAm_*/*Q_Oil_*= 0.08, ensuring the same TEMED amount availability across different *Q_PAAm_*/*Q_Oil_*. In this case, all droplets were suspended in the same oil volume. Our results confirmed that when the TEMED amount was varying, bead swelling decreased with increasing *Q_PAAm_*/*Q_Oil_* (Figure 5B). This indicated that higher TEMED concentration in the droplets, connected to lower *Q_PAAm_*/*Q_Oil_* (i.e., higher *V_Oil_*, since *V_PAAm_* is constant and equal to 50 µL), led to increased swelling. Conversely, when the TEMED amount was matched across all conditions (*Constant V_Oil_*), beads produced at different *Q_PAAm_*/*Q_Oil_* exhibited comparable swelling, confirming that TEMED concentration in droplets is a key parameter governing variations in microgel physical properties (Figure 5B).

**Figure 5.**
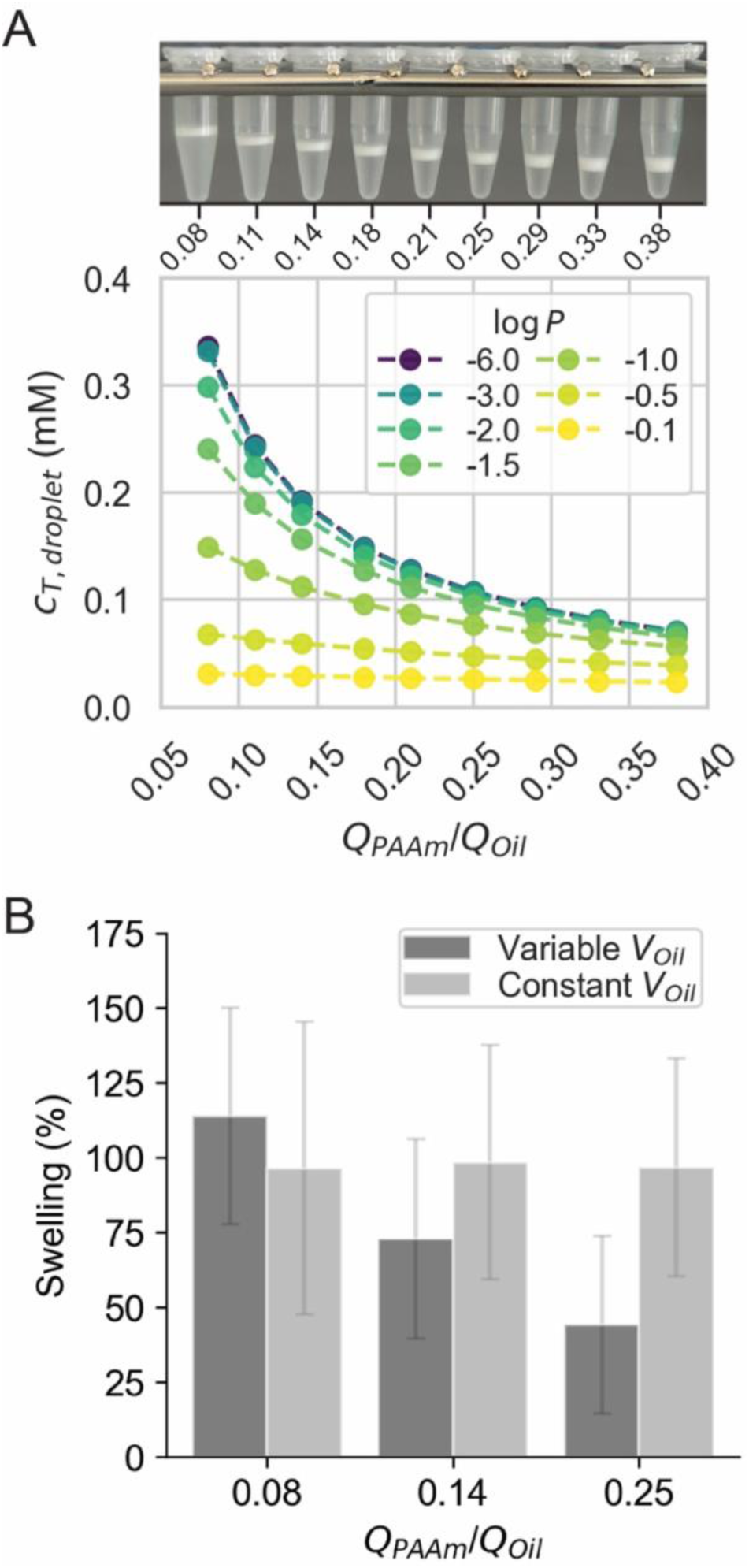
Flow-driven control of catalyst transport into droplets and its influence on bead swelling. Analysis of TEMED concentration in droplets (*c*_*T*,*droplet*_) as a function of *Q_PAAm_/Q_Oil_* ratio and its impact on microgel bead swelling. The droplets were generated at a constant total flow rate (*Q_Total_* = 20 µL/min), while varying *Q_PAAm_/Q_Oil_* ratio. **(A)** The graph shows possible values of *c*_*T*,*droplet*_, depending on the *Q_PAAm_/Q_Oil_* and the partition coefficient *P*, calculated using equation (8). Above the graph are pictures of the collection tubes containing pre-gel droplets produced at various *Q_PAAm_/Q_Oil_* ratios (0.08–0.38; corresponding values are reported below each vial). The white layer visible in the vials is the pre-gel droplets floating on top of the oil solution. The *Q_PAAm_/Q_Oil_* ratio directly modulates the volume ratio between pre-gel droplets and oil (*V_PAAm_*/*V_Oil_*). Since the *V_PAAm_* was kept constant (50 µL), only the *V_Oil_* varied. **(B)** Bar plot showing the swelling (%) of the final microgel beads in 1×PBS, polymerized from pre-gel droplets produced at three different *Q_PAAm_/Q_Oil_* ratios. Polymerization was performed under two conditions: *i*) Varying *V_Oil_* (dark gray bars), as determined by different *Q_PAAm_/Q_Oil_* ratios used during production (see vials in panel A). *ii*) Constant *V_Oil_* (light gray bars), where the oil volume in the collection tube was adjusted during production for the pre-gel droplets produced at *Q_PAAm_/Q_Oil_* = 0.14 and 0.25, so that the final oil volume matched that of the batch produced at *Q_PAAm_/Q_Oil_* = 0.08. The TEMED concentration in oil (*C_T,i_*), was kept constant across all conditions. Error bars represent propagated uncertainties derived from standard deviations of the bead and droplet volumes.

In particular, our results indicated that higher TEMED concentrations tend to inhibit gelation, since for lower values of the *Q_PAAm_*/*Q_Oil_*, the yield, mass conversion and Young’s modulus decreased drastically (Figure 3 and 4C). This behavior can be ascribed to the acceleration of radical initiation, increasing the rate of monomer radical formation, which reduces the average polymer chain length and results in less uniform crosslinking and incomplete polymerization, as reported before for the bulk gel [77,88,98]. Chrambach & Rodbard, observed a consistent decrease in extractable monomer with increasing TEMED concentration from ∼2 to 14 mM, suggesting an optimal TEMED concentration of 1–10 mM for reproducible and efficient polymerization [98]. Patel et al. showed that increasing the TEMED concentration from 0–260 mM reduced the mean elastic modulus of the PAAm gels from 1.9 kPa to 0.7 kPa [89]. Hara et al. reported a linear increase in the gel swelling with varying TEMED concentration between 1–5 mM [99].

It is well established that chemical reactions in microdroplets are typically faster and more uniform than in bulk systems. However, an important yet often overlooked aspect is the increased sensitivity of these microreactors. In our case, change in TEMED concentration, from 0.1 mM to 0.3 mM, corresponding to minor volume difference of only ∼50 fL, led to pronounced changes in the physical properties of the resulting microgels. These findings highlight the critical importance of precisely controlling the parameters that influence interfacial transport during microgel gelation, in order to ensure standardization and reproducibility in the fabrication of microgels with tailored physical properties.

Taken together, we have systematically shown for the first time that flow conditions, by modulating the transport of catalyst at the oil–water interface, can indirectly alter the physical properties of the microgel beads. Unlike bulk gels, in microfluidic polymerization, small changes in the reagent concentration result in measurable differences in the bead properties, even when pre-gel droplet diameter and monomer concentration are held identical. Notably, these findings introduce a simple yet effective strategy through which the microgel elasticity can be finely tuned by modulating the interfacial catalyst transport solely via flow control, without altering the pre-gel composition. This approach eliminates the need to prepare multiple pre-gel formulations separately, thereby reducing the risk of variability introduced by pipetting errors or inconsistencies in volume measurements. By preparing a single stock solution of acrylamide (AAm) and bis-acrylamide (BIS) in Tris buffer, which can be stored at 4°C, only the addition of APS prior to use is required. This streamlines the process and minimizes sources of error. This approach uniquely enables the production of beads with highly consistent sizes (diameter variation less than 1.2 µm) while varying their Young’s moduli (from 0.7 to 1.8 kPa), effectively decoupling size from elasticity, within a range of interest for different biophysical applications [7,17,18,26,28,50]. In contrast, standard production methods often require preparing and screening multiple batches, as adjusting the pre-gel composition to achieve a desired stiffness also affects the swelling behavior of the microgel meshwork, which in turn alters the final bead size. As we have shown, this process is further complicated by the system’s high sensitivity to even small variations in formulation or processing conditions. Nevertheless, we also identified a range of the flow rate ratio (0.21 ≤ *Q_PAAm_*/*Q_Oil_* ≤ 0.29) for a fixed value of the total flow rate (*Q_Total_* = 20 µL/min) in which the properties of the final beads were less sensitive to flow variations. Within this range, we observed more consistent and efficient production, with increased droplet frequency, a higher polymerization yield and reduced variability in bead size and elasticity among different flow rate ratios.

Collectively, our findings deliver a comprehensive and compelling analysis of critical parameters for the standardized, reliable and reproducible production of cell-mimicking microgel beads, with tailored size and elasticity of interest for several biophysical applications. Furthermore, we identify the optimal conditions to enable high-throughput, efficient fabrication of PAAm microgel beads, paving the way for scalable applications.

## Conclusion

In this study, we have demonstrated that flow conditions during droplet production critically influence the physical properties of PAAm microgel beads, particularly their elasticity, even when pre-gel composition and droplet diameter are identical. Using a flow-focusing microfluidic geometry, we identified a droplet production regime in which varying the PAAm-to-oil flow rate ratio at a constant total flow rate yielded monodisperse droplets with < 1 µm diameter variation with identical pre-gel composition. This allowed us to systematically investigate the effect of flow conditions on polymerization and physical properties of the resulting microgel beads, independent of the droplet diameter and pre-gel composition. We found that increasing the flow rate ratio led to: (i) higher polymerization yield, (ii) reduced bead swelling, and (iii) increased Young’s modulus. We identified the concentration of catalyst within droplets, modulated by the flow rate ratio, as a key factor driving variations in the final microgel properties. These findings highlight the extreme sensitivity of microscale in-drop polymerization to interfacial transport phenomena at the oil–water interface. While these effects are negligible in bulk gels, they become critical in microreactors, which are highly susceptible to minor fluctuations in chemical concentration.

Our results highlight a crucial yet often overlooked factor in microgel production, providing insights into microfluidic parameters needed for the standardized, reproducible, and scalable fabrication of PAAm microgels. Together, this work paves the way for high-throughput production of PAAm microgel beads with precisely tunable physical properties, pivotal for developing novel biomaterials that advance applications in mechanobiology, diagnostic standardization, tissue engineering and drug delivery.

Our results also define the optimal conditions for high-throughput and efficient fabrication of PAAm microgel beads, which are clearly translated into the distribution and movement of droplets immediately after their formation at the crossjunction, leading to the emergence of characteristic droplet stream patterns. Looking ahead, these visual data could be used to train machine learning algorithms for real-time monitoring and automatic adjustment of flow conditions, enabling the precise production of microgels with desired physical properties. Additionally, differences in refractive index closely linked to properties such as Young’s modulus suggest that image-based analysis could allow rapid, non-invasive characterization of bead mechanical properties. Integrating these capabilities into a microfluidic platform would enable on-demand production and in-line characterization of tailored microgel beads, greatly advancing their applicability in research and industry.

## Materials and Methods

### Microfluidic chip design and master mold fabrication

The microfluidic device design featuring flow-focusing geometry was designed using KLayout software. The device design included two inlets, each for polyacrylamide pre-gel (dispersed phase) and oil solution (continuous phase) and one outlet to collect the pre-gel droplets. An array of circular pillars (diameter = 100 µm with a gap ranging from 15–100 µm) was included after the inlet chambers to avoid clogging in the channels. A flow-focusing junction named a ‘crossjunction’ was designed such that the inlet channels for both solutions meet at 90° angle with a channel width of 15 µm. Post-crossjunction channel gradually widens to 250 µm further ending at the outlet chamber. The master mold was fabricated via a photolithography process (SUSS MA/BA6 Gen4 Semi-automated mask aligner) using AZ15nXT 450 cps photoresist (MicroChemicals GmbH). A 15 µm thick layer of phototresist was spin-coated at 2000 rpm, and 5000 rpms/s acceleration for 9.5 s (Laurell WS-650Hzb-23NPP-UD-3 spin coater) on a 4-inch silicon wafer. The coated wafer was soft baked at 110 °C for 3 min, followed by a UV light exposure of 450 mJ/cm^2^ through a chrome photomask containing device designs. Post-exposure, the wafer was baked at 120 °C for 1 min and was developed in a solution of AZ 400K developer (1:3 v/v, developer:distilled water) for 3 minutes and 10 seconds, to realize the device structures on the master mold. The height of the resulting microstructures was measured using an optical profilometer (KLA Zeta™-20). Prior to use, the master template was functionalized with 1H, 1H, 2H, 2H-perfluorodecyltriethoxysilane (Sigma-Aldrich) in a desiccator for 12 h to render its surface hydrophobic for an easy peel-off of the PDMS replica.

### Microfluidic chip fabrication

Polydimethylsiloxane (PDMS) microfluidic chips were fabricated by replicating the master features using a soft lithography process. The PDMS base (Dow Corning Sylgard® 184) and curing agent were mixed in a 10:1 w/w ratio, degassed, poured over the master, and polymerized in an oven at 75 °C for 1 h 15 min. Once cured, the PDMS replica was carefully peeled off from the master and punched at the inlets (I.D. 1.5 mm) and outlet (I.D. 1.5 mm) chambers using a biopsy puncher (Imtegra GmbH). The punched PDMS structure was then bonded to a glass coverslip (40 × 24 mm^2^, thickness 2, Hecht, Germany) by surface activation using an air plasma treatment (Plasma system Atto, Diener Electronic, Germany) at 75 W for 2 min. Finally, the assembled PDMS chip was incubated at 75 °C for 12 h to ensure stable bonding.

### Surfactant synthesis

The preparation of the ammonium carboxylate salt of Krytox® was carried out using a slightly modified protocol from the literature [100]. To summarize, Krytox® 157 FSH (10 g, DuPont) was dissolved in a solvent mixture of 60 mL methanol (Sigma Aldrich) and 30 mL of 3M™ Novec™ 7100 (IoLiTec Ionic Liquids Technologies GmbH) under stirring until completely dissolved. Subsequently, 25 mL of 0.1 N ammonium hydroxide (Sigma Aldrich) was added dropwise while maintaining vigorous stirring. The reaction proceeded overnight under continuous stirring. After the reaction, the solvent was removed under reduced pressure and the product was dried under high vacuum to yield a pale, viscous oil.

### Polyacrylamide microgel bead fabrication

To facilitate the stable production of polyacrylamide (PAAm) pre-gel droplets in oil and to keep the aqueous phase from wetting the microfluidic channel, the inner walls of the chip were functionalized by flowing Aquapel® (Pittsburgh Glass Works). The excess of Aquapel® was flushed out by pressurized air and the device was allowed to dry at 65°C for 5 min. The oil solution was prepared by mixing 2% w/w of ammonium Krytox® surfactant and 0.4% v/v N, N, N′, N′-tetramethylethylenediamine (TEMED) (Sigma Aldrich) in 3M™ Novec™ 7500 (IoLiTec Ionic Liquids Technologies GmbH). The PAAm pre-gel solution was prepared by diluting 40% acrylamide (Sigma Aldrich), 40% acrylamide/bisacrylamide 19:1 (Sigma Aldrich) and 10% w/v ammonium persulfate (GE Healthcare) in 10 mM Tris buffer (pH 7.5) such that the total monomer concentration (*C_T_*) was 5.6% w/v. The flow of both solutions was initiated by pressurizing the fluid-containing vials until both solutions entered the microfluidic channels. Once the solutions filled the channels and no air bubbles were trapped in the fluidic path, specific flow rates for each of the solutions were applied to obtain *Q_PAAm_*/*Q_Oil_* as described in Figures 2A and 3A. As the flow rates were established and stable droplet production was achieved, produced pre-gel droplets were collected until 50 µL of the PAAm pre-gel solution was consumed for each of the *Q_PAAm_*/*Q_Oil_*. The collected pre-gel droplets were polymerized overnight at 65°C. The polymerized microgel beads in oil were washed three times with each of 20% v/v 1H, 1H, 2H, 2H-Perfluoro-1-octanol (Sigma Aldrich) in 3M™ Novec™ 7500, 1% v/v Span® 80 (Sigma Aldrich) in n-hexane (Sigma Aldrich) and 1×PBS (without calcium and magnesium, pH 7.4) (Gibco™) solutions via centrifugation at 5000×g for 5 min. The final bead suspension in 1×PBS was stored at 4°C.

### Polyacrylamide pre-gel droplet and microgel bead characterization

#### Droplet diameter & production frequency

To analyze droplet size distribution, bright-field images of pre-gel droplets were acquired during production in the post-crossjunction channel using an inverted microscope (Zeiss AxioObserver.A1) equipped with a 20×/0.45 objective. The acquired bright-field images were analyzed using a custom macro implemented in the open-source software Fiji to calculate the size distribution. The droplet production frequency was calculated as 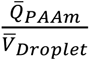, where *Q̄*_*PAAm*_ is the mean flow rate of PAAm pre-gel solution and *V̄*_*Droplet*_, the mean droplet volume was calculated from the droplet diameter. The number of droplets produced (*N*_*Droplet*_) was estimated from (*V*_*PAAm*_⁄*V̄*_*Droplet*_), where *V*_*PAAm*_ is the volume of PAAm pre-gel solution used to produce droplets (= 50 µL).

#### Bead yield and diameter

To estimate the yield, first, the number of beads polymerized (*N*_*Bead*_) was measured using a Neubauer haemocytometry chamber using brightfield microscopy (Zeiss AxioObserver.A1, Plan-Apochromatic 10×/0.3). The yield (%) was calculated as (*N*_*Bead*_ ⁄*N*_*Droplet*_) · 100. To assess microgel bead size distribution, bright-field images of sedimented beads in 1×PBS were acquired using the same microscope and a Plan-Apochromatic 10×/0.3 objective. The acquired bright-field images were analyzed using a custom macro implemented in the open-source software Fiji to calculate the size distribution.

### Optical Diffraction Tomography

To ensure optimal light transmission through the sample, a low-concentration bead solution was placed between two coverslips, held parallel to each other using double-sided tape as a spacer. The optical diffraction tomography (ODT) imaging was performed using a custom-built interferometric setup based on a Mach-Zehnder configuration, which enabled the acquisition of complex optical fields from multiple illumination angles, following previously established methodologies [72]. In this setup, a continuous-wave laser (λ = 532 nm, CNI Optoelectronics Technology Co.) was split into two beams using a beam splitter. One beam served as the reference, while the other was directed to illuminate the sample mounted on an inverted microscope (Axio Observer 7, Carl Zeiss AG) through a tube lens (f = 175 mm) and a water-dipping objective lens (40×, NA = 1.0, Carl Zeiss AG). Angular scanning of the illumination was achieved by a dual-axis galvanometric mirror (Thorlabs, GVS212/M) positioned at a plane optically conjugate to the sample, covering 150 incident angles for full tomographic acquisition. The light scattered by the specimen was collected by a second high-NA objective lens (63×, NA = 1.3, oil immersion, Carl Zeiss AG) and interfered with the reference beam to produce a hologram at the image plane. This hologram was recorded using a CMOS camera (MQ042MG-CM-TG, XIMEA). Complex field information was retrieved from the holograms using a Fourier transform-based reconstruction algorithm [101]. The three-dimensional refractive index distribution was then calculated using the retrieved complex fields, applying the Fourier diffraction theorem with the Rytov approximation [102,103]. Further methodological details are available in earlier publications [104], and the reconstruction scripts in MATLAB are publicly accessible at https://github.com/OpticalDiffractionTomography/ODT_Reconstruction (v1.0.0).

From the reconstructed RI tomograms, the dry mass of beads, *m*_*Bead*_, was calculated by integrating the RI contrast within the segmented volume, as 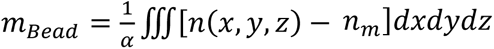, where *α* is the RI increment (dn/dc) with *α* = 0.19 mL/g for polyacrylamide [105] and *n*_m_ is the RI value of surrounding PBS, measured to be 1.337 using an Abbe refractometer (ORT1RS, Kern & Sohn GmbH).

### Real-time Deformability Cytometry (RT-DC)

RT-DC measurements were conducted using an AcCellerator device (Zellmechanik Dresden GmbH, Germany). PAAm microgel beads, suspended in 0.6% w/w MC-PBS buffer [106], at a concentration of 3–5 × 10⁶ beads/mL, were introduced into FEP tubing (I.D. 750 µm, Postnova Analytics GmbH) and then flowed into a PDMS-based chip with 30 µm square channels. The total flow rate was set to 0.12 µL/s using a syringe pump (neMESyS 290 N, Cetoni GmbH). All measurements were performed at a controlled chamber temperature of 25 °C. Imaging was carried out on an inverted microscope (Axio Observer Z1, Zeiss, Germany) with a 40× objective (EC-Plan-Neofluar, 40×/0.75; no. 420360-9900, Zeiss). Data acquisition and real-time contour analysis were performed using ShapeIn2 software (Zellmechanik Dresden). Data analysis was carried out with the open-source ShapeOut2 software [107], where the Young’s moduli of the beads were determined using a Buyukurganci viscosity model and HE-3D-FEM-22 lookup table after filtering events within a porosity range of 1.00–1.05 [84,106].

### Statistical analysis

All relevant statistical details are reported in the respective figure captions. The data shown in the main figures are from a representative experiment, which was independently repeated at least three times with similar results (Figure S10). Figure S10 summarizes droplet diameter, bead diameter and Young’s modulus data from three independent experimental replicates.

## Data availability

The data published in this study are available from the authors upon request.

## Author contributions

Conceptualization: R.G., S.G.; Investigation: R.G.; Methodology: R.G., S.G., K.K.; Data curation: R.G., K.K., S.G.; Validation: R.G.; Visualization: R.G., S.G.; Writing (original draft): R.G., S.G.; Writing (editing and review): all authors; Funding acquisition: S.G., J.G.; Supervision: S.G., J.G., A.R.B.

## Acknowledgments

We thank Parth Patel and Mahi Muraleedharan, part of the Core Lab-on-a-chip Facility at Max-Planck-Zentrum für Physik und Medizin, for the production of the microfluidic chips used for the bead production and RT-DC measurements. We are grateful to Cornelia Liebers for preparing 0.6% MC-PBS solution. We thank Catherine Xu and Sandra García Rey for valuable discussions during this study. We thank the TDSU Micro-& Nanostructuring facility at Max Planck Institute for the Science of Light for providing access to the cleanroom infrastructure and well-maintained microfabrication instruments. We also thank Ralf Keding for his assistance with surfactant synthesis. We thank ChatGPT (OpenAI) for assistance with language editing and refinement of the manuscript. The authors acknowledge the financial support by the European Union’s Horizon 2020 research and innovation program under grant agreement No. 953121 (project FLAMIN-GO) and the funding from the Max Planck Society to J.G.

