## Supplementary Data for "Fine-tuning cell-mimicking polyacrylamide microgels: Sensitivity to microscale reaction conditions in droplet microfluidics"

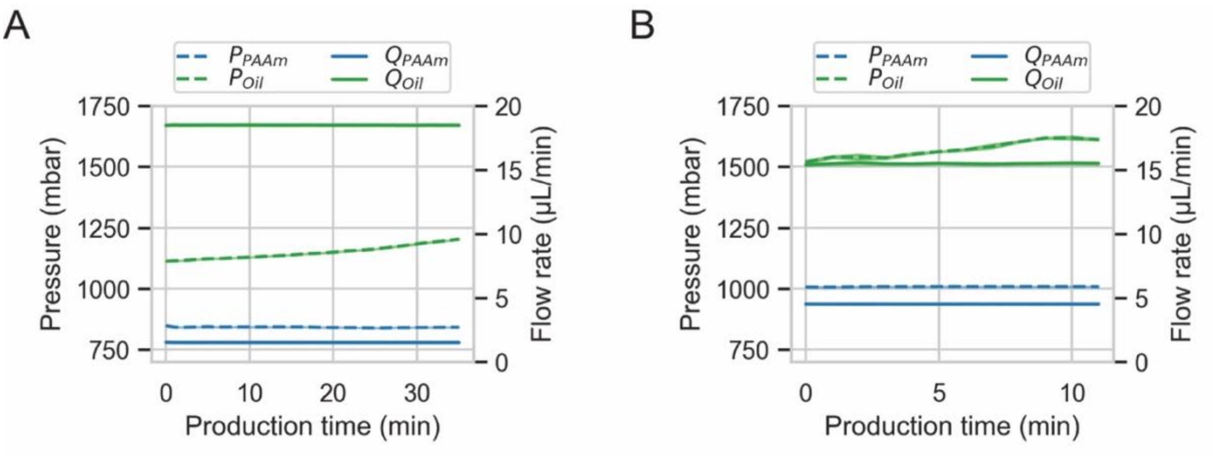


**Figure S1 Closed-loop feedback control during PAAm pre-gel droplet production.** Automatic pressure adjustments of PAAm pre-gel ($P_{PAAm}$) and oil ($P_{Oil}$) solutions to maintain constant flow rates of PAAm ($Q_{PAAm}$) and oil ($Q_{Oil}$) over the entire production time. The graphs show these parameters over time during two representative droplet production experiments. The targeted flow rates were (A) $Q_{PAAm}$ (blue solid line) = 1.5 µL/min and $Q_{Oil}$ (green solid line) = 18.5 µL/min, and (B) $Q_{PAAm}$ = 4.5 µL/min and $Q_{Oil}$ = 15.5 µL/min. For each experiment, the droplet production was performed using a fixed volume (50 µL) of PAAm pre-gel solution. The lines represent the mean values, while the shaded bands indicate standard deviations across all data points within each 1-min bin.

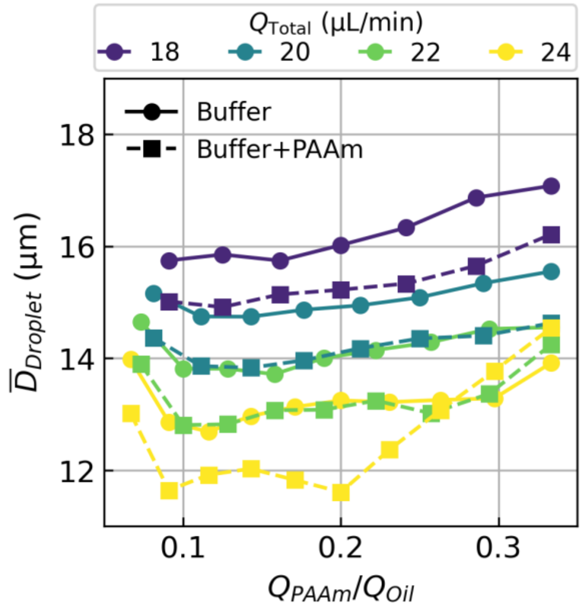


**Figure S2 Effect of dispersed phase composition on droplet size.** Mean droplet diameter as a function of $Q_{PAAm}/Q_{Oil}$ ratio at different total flow rates ($Q_{Total}$), using Tris buffer (●) and Tris buffer containing PAAm pre-gel components (■) as the dispersed phase at $C_T$ = 5.6% w/v.

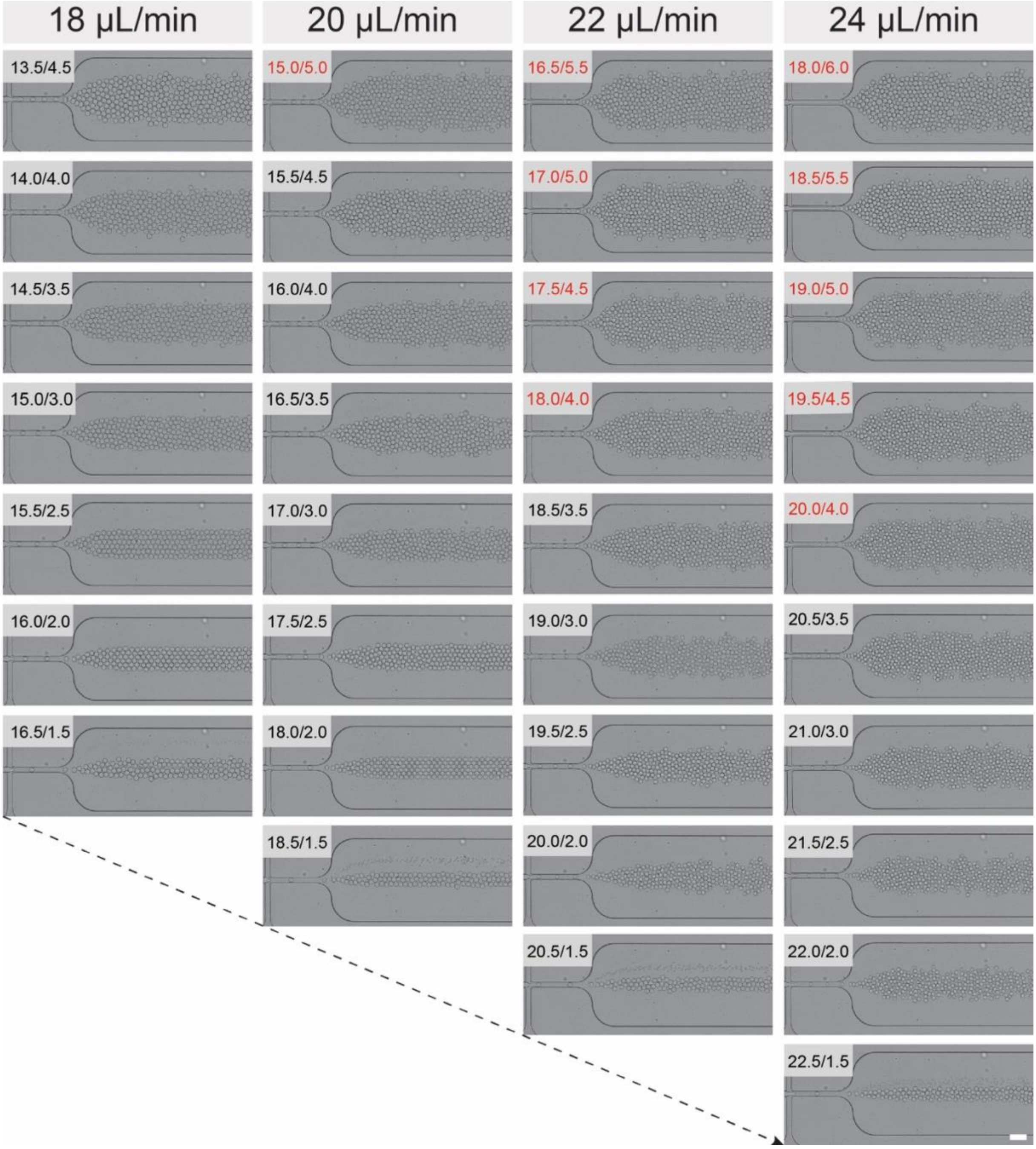


**Figure S3 Brightfield images showing droplet generation at the flow-focusing crossjunction under varying flow conditions.** Images are grouped by $Q_{Total}$ (18, 20, 22 and 24 µL/min), with corresponding combinations of $Q_{Oil}$ and $Q_{PAAm}$ indicated as $Q_{Oil}/Q_{PAAm}$ (in black for dripping and in red for jetting regime) on each image. Scale bar is 50 µm. Images along the diagonal (see arrow) show droplet formation at a fixed $Q_{PAAm}$ and increasing $Q_{Oil}$.

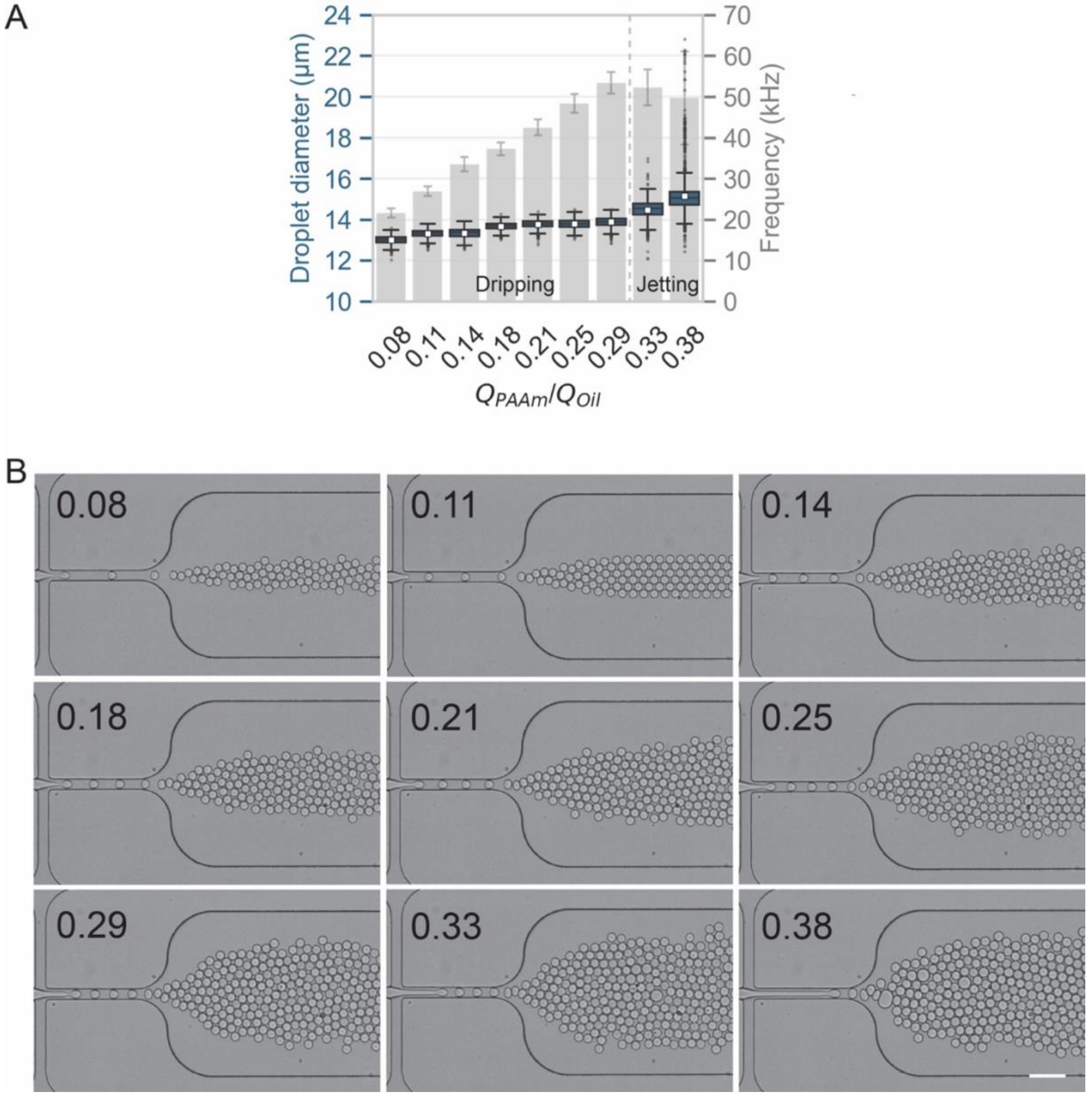


**Figure S4 Effect of $Q_{PAAm}/Q_{Oil}$ ratio on pre-gel droplet production.** Droplet diameter and production frequency as a function of $Q_{PAAm}/Q_{Oil}$ for a fixed $Q_{Total}$ = 20 µL/min. **(A)** The box plots represent the droplet diameter distribution at varying $Q_{PAAm}/Q_{Oil}$. Each box represents the interquartile range (IQR) covering data between 25th and 75th percentiles. The horizontal line within the box represents median, while the white square markers indicate mean. Whiskers extend to 1.5 times the IQR, while the data points outside this range are represented as dots, which indicate outliers. The bar plot depicts the droplet production frequency (kHz) as a function of $Q_{PAAm}/Q_{Oil}$. The frequency was calculated as the ratio of $\bar{Q}_{PAAm}$ and mean droplet volume, $\bar{V}_{Droplet}$, while the error bars represent the propagated uncertainty based on the standard deviation of the $\bar{Q}_{PAAm}$ and $\bar{V}_{Droplet}$. The dashed line separates the two droplet production regimes: dripping and jetting. **(B)** Bright-field images showing pre-gel droplet formation at the crossjunction for different $Q_{PAAm}/Q_{Oil}$ values (indicated in the top-left corner of each image), corresponding to the data presented in (A). Scale bar: 50 µm.

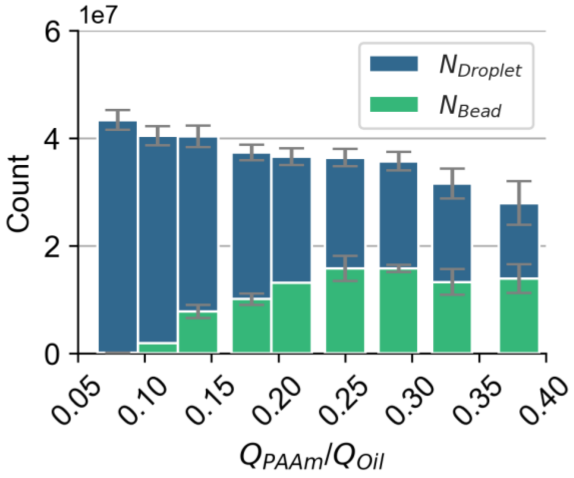


**Figure S5 Comparison of the number of droplets and beads as a function of $Q_{PAAm}/Q_{Oil}$.** Analysis of the number of droplets produced and the number of beads polymerized. The bar plots show the estimated number of produced droplets ($N_{Droplet}$, in blue) and the measured number of polymerized beads ($N_{Bead}$, in green). $N_{Droplet}$ was calculated as the ratio of the volume of PAAm used during the entire droplet production (50 µL) to the droplet volume. Error bars represent the standard deviations.

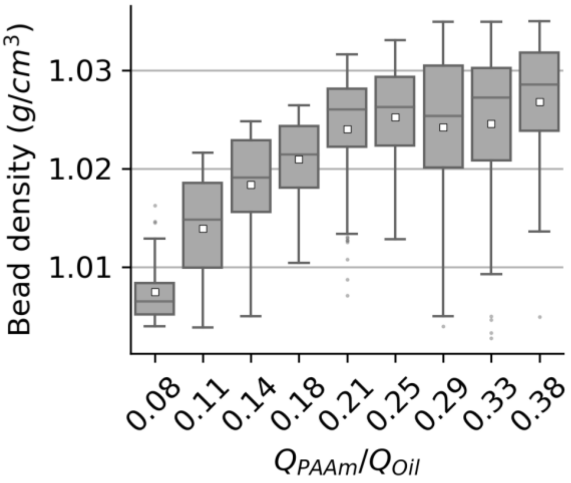


**Figure S6 Density of PAAm beads as a function of $Q_{PAAm}/Q_{Oil}$.** Analysis of bead density obtained from ODT measurements. Box plots showing the distribution of bead density (g/cm$^3$) of individual beads produced at varying $Q_{PAAm}/Q_{Oil}$. Each box represents the interquartile range (IQR) covering data between 25$^{th}$ and 75$^{th}$ percentiles. The horizontal line within the box represents median, while the white square markers indicate mean. Whiskers extend to 1.5 times the IQR, while the data points outside this range are represented as dots, which indicate outliers.

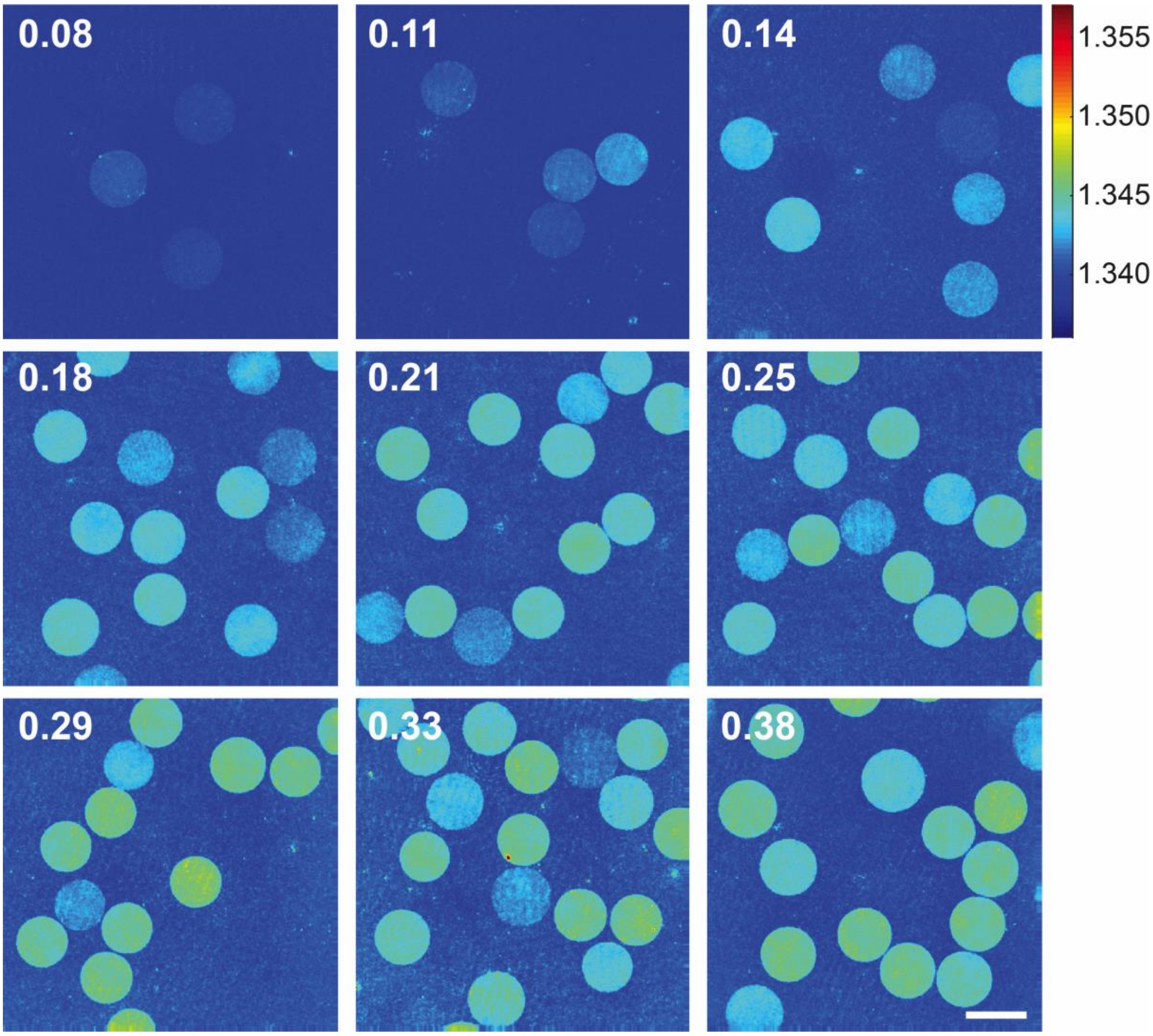


**Figure S7 Refractive index analysis of PAAm beads using ODT.** Refractive index tomograms of the PAAm microgel beads suspended in 1×PBS, acquired using ODT. Beads were obtained from the pre-gel droplet produced at varying $Q_{PAAm}/Q_{Oil}$ values, as depicted above each tomogram. The color scalebar indicates refractive index values, ranging from 1.340 to 1.355. The scale bar is 20 µm.

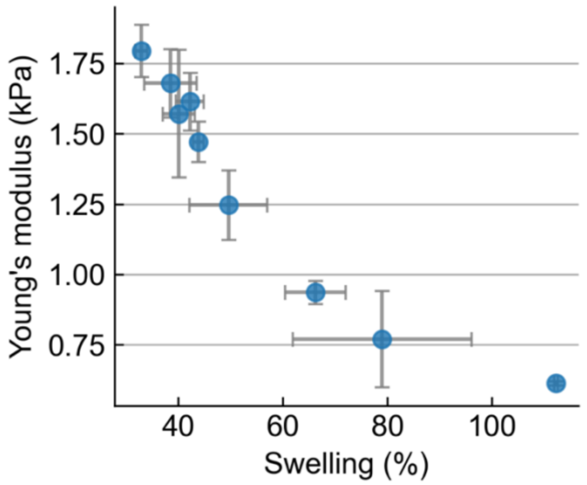


**Figure S8 Relationship between the swelling behavior and elasticity of the PAAm microgel beads.** A scatter plot showing mean values of the swelling (%) and Young's modulus of the PAAm microgel beads obtained from three independent experiments. Swelling (%) was calculated according to equation (4). The Young's modulus was analyzed using RT-DC as described in Figure 4. The error bars represent the standard deviations.

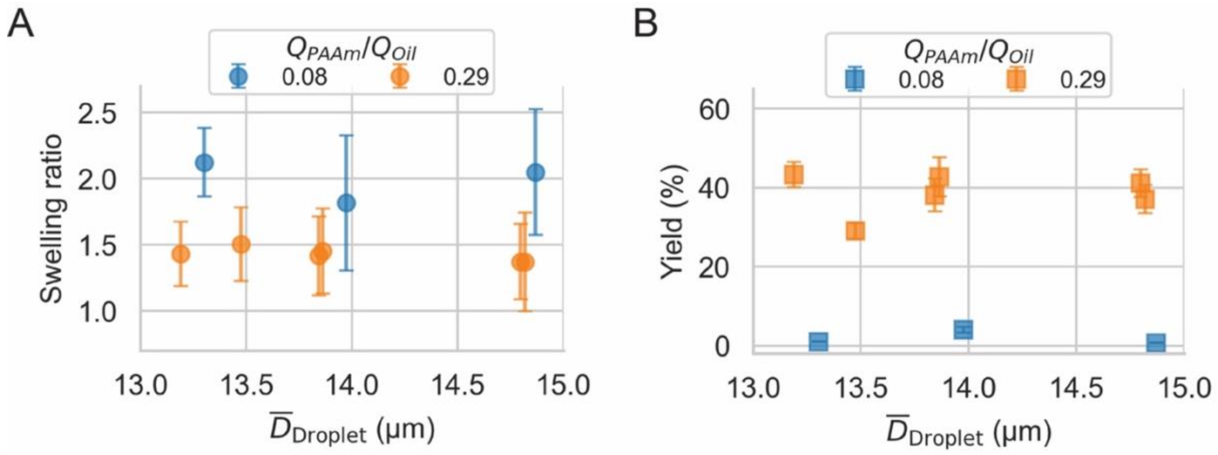


**Figure S9 Relationship between pre-gel droplet diameter and the swelling behavior and yield of PAAm microgel beads.** A scatter plot showing mean values of the **(A)** swelling (%) and **(B)** yield (%) of the PAAm microgel beads vs their corresponding pre-gel droplet diameters produced under two different $Q_{PAAm}/Q_{Oil}$ ratios. Yield (%) and Swelling (%) were calculated according to equations (3) and (4), respectively. The error bars for swelling represent the propagated uncertainty calculated using the standard deviations of bead and droplet diameters. The error bars for yield represent the propagated uncertainty derived from the standard deviations of $N_{Droplet}$ and $N_{Bead}$.

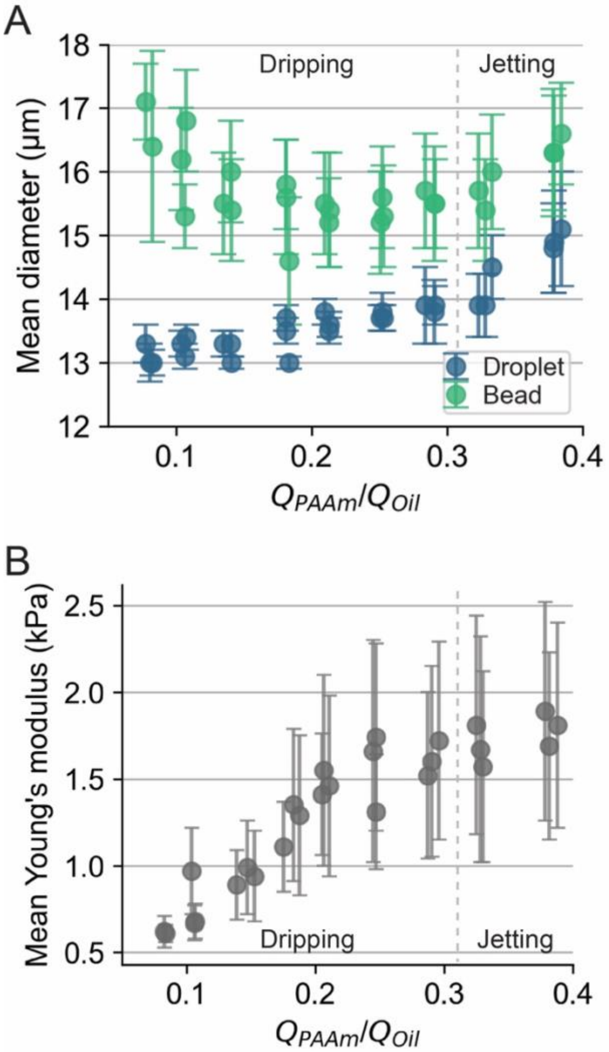


**Figure S10 Experimental repetitions showing reproducibility and variation in droplet diameter, bead diameter and bead Young's modulus as a function of $Q_{PAAm}/Q_{Oil}$.** Analysis of the PAAm microgel bead diameter and elasticity obtained from the polymerization of pre-gel droplets produced at a constant $Q_{Total}$ = 20 µL/min and varying $Q_{PAAm}/Q_{Oil}$. **(A)** Comparison between mean droplet diameter (blue) and mean bead diameter (green) at varying $Q_{PAAm}/Q_{Oil}$ ratios for three independent experiments for each condition. For the condition $Q_{PAAm}/Q_{Oil}$ = 0.08, bead polymerization failed in one of the replicates. The error bars represent the standard deviation. **(B)** Mean Young's modulus of the PAAm microgel beads analyzed by RT-DC as a function of $Q_{PAAm}/Q_{Oil}$ for three independent experiments. The error bars represent the standard deviations. The dashed line separates the two droplet production regimes: dripping and jetting.
